# Hierarchical buffering of cellular RNA polymerase II pools maintains transcriptional homeostasis

**DOI:** 10.64898/2025.12.22.695869

**Authors:** Alexander Gillis, Roberta Cacioppo, Pei Qin Ng, Hongpei Li, Bailey Hiles-Murison, Ana Tufegdžić Vidaković, Scott Berry

## Abstract

RNA polymerase II (Pol II) abundance and transcriptional output are coordinated across diverse conditions to maintain mRNA homeostasis. To determine whether Pol II acts as a limiting factor for transcription, we combined acute perturbations with quantitative single-cell imaging and genome-wide profiling. We find that transcription remains stable across 70-180% of normal Pol II levels in human cells. This robustness emerges from hierarchical buffering mechanisms operating on distinct timescales. Within tens of minutes after monoallelic depletion of Pol II subunit POLR2A, cells maintain steady transcriptional output by drawing polymerases from a nucleoplasmic reserve pool. However, over hours, cellular Pol II holoenzyme levels also recover, driven by constitutive excess POLR2A subunit production rather than feedback sensing. Finally, strong depletion over days activates additional compensatory mechanisms. Together, these results reveal how temporal separation of buffering mechanisms provides robustness to perturbations of essential machinery, illuminating how cells maintain homeostatic control of essential complexes.

## Introduction

Concentration homeostasis is fundamental to cellular function. Maintaining constant concentrations of mRNA and protein is underpinned by coordinating transcription with both cell size and mRNA degradation (*1, 2*). RNA polymerase II (Pol II), the twelve-subunit enzyme complex responsible for transcribing all mRNA, plays a central role in this (*3*). Pol II abundance and global transcriptional output change in parallel across diverse cellular conditions, suggesting tight regulatory coupling (*4, 5*). However, whether Pol II abundance limits transcriptional capacity – directly determining output – or whether homeostatic mechanisms actively buffer the fluctuations, perturbations, and functional modulations that occur during normal cellular processes, has remained unclear. In yeast, evidence supports a limiting-factor model in which transcription is determined directly by Pol II levels (*5*). Whether human cells operate under similar constraints remains unknown, despite the broader implications for understanding how cells maintain concentration homeostasis for essential, stoichiometrically assembled complexes.

Several features of the transcriptional machinery could buffer changes in Pol II abundance. First, cellular Pol II distributes between chromatin-bound and nucleoplasmic pools, with approximately half engaged in transcription at any time (*3*). Thus, redistribution from the nucleoplasmic pool could buffer changes in total Pol II abundance that occur during cell growth, cell cycle progression, or due to stochastic variation in subunit levels. Second, multiple checkpoints control polymerase usage efficiency (*6, 7*). Pol II molecules bound at promoters must escape from preinitiation complexes through TFIIH-mediated phosphorylation of the repetitive C-terminal domain (CTD), and at many genes, there is also pausing downstream of transcription start sites, where polymerases either terminate or are released into productive elongation – dependent on positive transcription elongation factor b (P-TEFb) (*8*). If Pol II becomes limiting, modulating these checkpoints could maintain cellular transcriptional output, by increasing output per polymerase. Third, Pol II abundance itself could be adjusted through altered synthesis or stability of its twelve stoichiometric subunits, or through complex assembly (*9, 10*). Because these mechanisms would operate on different timescales – minutes for redistribution versus hours for abundance changes – multiple hierarchical buffering mechanisms could provide robustness against variation in Pol II levels.

However, distinguishing these mechanisms experimentally requires approaches that avoid confounding effects while enabling temporal resolution. Complete depletion of essential proteins like Pol II triggers cell death and pleiotropic stress responses (*11*–*13*) that obscure homeostatic mechanisms, while gradual depletion risks conflating direct responses with secondary adaptations. To address these challenges, we developed a monoallelic depletion strategy that acutely removes POLR2A, the largest Pol II subunit, from one allele while leaving the other intact for monitoring. This approach reduces total Pol II abundance while maintaining cell viability and enabling quantitative measurements of the unperturbed polymerases. Using these cells, we tracked Pol II transcriptional activity, chromatin-bound and nucleoplasmic pools using quantitative single-cell time-lapse microscopy and genome-wide profiling, with measurements spanning minutes to days following perturbation.

## Results

### Pol II abundance does not limit transcription in human cells

To examine the extent to which Pol II abundance determines transcriptional output in human cells, we engineered inducible miniAID-mClover degron tags into both alleles of endogenous POLR2A in HCT116 cells (*12*) (Fig. 1A-B, Fig. S1). If Pol II were a limiting factor for transcription, decreases in Pol II abundance should produce proportional decreases in transcriptional output. Treating mAC-POLR2A degron cells with high concentrations of auxin analog 5-phenyl indole-3 acetic acid (5-Ph-IAA; 100 nM - 10 µM) (*14*) drove rapid Pol II depletion. However, elongating Pol II marked by POLR2A CTD phosphorylation at Serine 2 (pSer2, measured using immunofluorescence) was only modestly reduced (Fig. 1B-D, Fig. S2A-B). Lower concentrations of 5-Ph-IAA (10 nM) which reduced total POLR2A abundance by 30 ± 5% (mean ± S.D.), also did not induce a proportional decrease in pSer2 levels (6 ± 5% reduction) (Fig. 1B,E-F, S2B). These results indicate that Pol II is present in excess rather than maintained at levels that limit cellular transcription.

**Figure 1.**
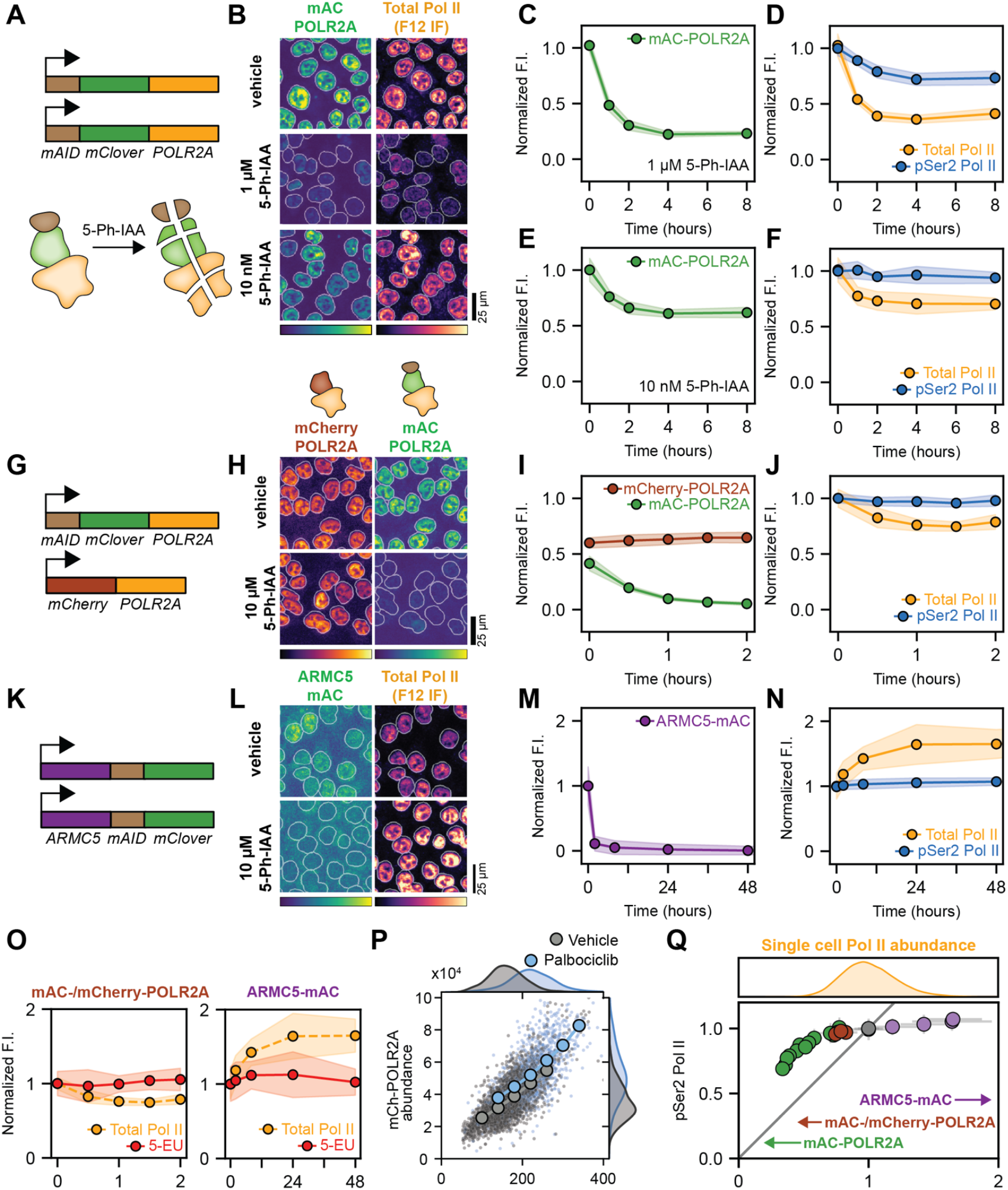
Pol II activity is robust to manipulation of Pol II abundance. (**A**) Biallelic degron tagging of POLR2A. (**B**) Imaging of mClover fluorescence and total Pol II (immunofluorescence) in mAC-POLR2A cells at high (1 µM) and low (10 nM) 5-Ph-IAA concentrations (4 h). Nuclear segmentation used for quantification shown in white outlines. (**C**) mClover fluorescence in mAC-POLR2A cells after 1 µM 5-Ph-IAA addition, normalised to unperturbed levels. (**D**) Total Pol II (F12) and pSer2 Pol II (3E10) immunofluorescence. (**E**,**F**) as in C,D, respectively, for 10 nM auxin. (**G**) Tagging one POLR2A allele with mAID-mClover and the other allele with mCherry. (**H**) Imaging of mClover and mCherry fluorescence in vehicle and 5-Ph-IAA-treated cells (2 h). (**I**) mClover and mCherry fluorescence in monoallelic mAC-POLR2A / mCherry-POLR2A cells after 10 µM 5-Ph-IAA addition, normalised to fluorescence levels in biallelic mAC-POLR2A cells and biallelic mCherry-POLR2A cells, respectively. (**J**) as in D, for monoallelic degron cells. (**K**) Schematic of ARMC5-mAID-mClover degron tagging. (**L**) Imaging of mClover fluorescence and total Pol II (immunofluorescence) in ARMC5-mAC cells in vehicle and 5-Ph-IAA treated cells (48 h). (**M**,**N**) As in C,D, for ARMC5-mAC cells. (**O**) 5-EU incorporation during 5-Ph-IAA treatment of monoallelic POLR2A degron and ARMC5-mAC cells.(**P**) Abundance (integrated nuclear fluorescence intensity) of mCherry-POLR2A compared to nucleus area following vehicle or palbociclib treatment. (**Q**) Summary of data from D,F,J, and N showing relationship between Pol II activity (pSer2 Pol II) and Pol II abundance (F12). Line indicates proportionality. Distribution of total Pol II in unperturbed cells from P shown for comparison. Note some error bars do not extent beyond points and are not visible. All fixed-cell immunofluorescence and 5-EU data is shown as mean ± S.D. of replicate wells collected in 2-3 separate experiments.

A potential confound of this biallelic degron approach is that ubiquitin-mediated degradation directly targets the same Pol II complexes being measured, which may alter Pol II activity independently from abundance changes. To circumvent this, we generated cells in which only one POLR2A allele carries the miniAID-mClover degron tag (mAC) while the other is tagged with the distinguishable fluorophore mCherry (mAC-/mCherry-POLR2A cells) (Fig. 1G-H, Fig. S1). This allows depletion of one pool of Pol II while monitoring the other, unperturbed pool. In monoallelic degron cells, high concentrations of 5-Ph-IAA led to loss of mAC-POLR2A with a half-life of ∼20 minutes while leaving mCherry-POLR2A levels unchanged (108 ± 7%) over 2 hours (Fig. 1I). This reduced total Pol II to 75 ± 4% of baseline at 90 minutes yet elongating Pol II remained stable, as quantified by pSer2 immunofluorescence (Fig. 1J). Separation of depleted and measured Pol II pools demonstrates transcriptional robustness to reduced polymerase abundance without artifacts from directly targeting the active complexes.

To increase Pol II levels beyond those in unperturbed cells, we generated cells with an inducible degron tag on ARMC5, key component of a ubiquitin ligase that normally constrains POLR2A levels during homeostasis (*15*–*17*) (Fig. 1K-L, Fig. S3). 5-Ph-IAA-mediated degradation of ARMC5-miniAID-mClover gradually elevated total Pol II by close to two-fold over 2 days, however pSer2 levels remained stable (Fig. 1M-N). This again contradicts a limiting factor model, indicating Pol II is already in excess at normal levels.

As an independent assay of cellular transcription, we quantified the incorporation of 5-ethynyl uridine (5-EU) into nascent RNA at the single-cell level via click chemistry (*4, 18*). During 5-Ph-IAA treatment of either monoallelic POLR2A or ARMC5 degron cells, where Pol II levels were either reduced to 75% or increased to 180% of baseline, 5-EU incorporation remained stable (Fig. 1O), confirming that transcription is buffered against these abundance changes.

Pol II abundance and nuclear volume scale with cell volume (*4, 19*), maintaining constant Pol II nuclear concentration. This Pol II abundance scaling is maintained even when cell size is increased beyond its normal range with CDK4/6-inhibitor, palbociclib (Fig. 1P, Fig. S4A). Despite this average scaling, we observe considerable cell-to-cell variability in Pol II concentration (*4*) (Fig. 1Q). This raised the question whether natural variability in Pol II levels might challenge transcriptional homeostasis. Combining results from all Pol II modulations tested revealed that transcription remains stable across 70-180% of average unperturbed Pol II levels (Fig. 1Q), demonstrating substantial robustness. Only below ∼70% of baseline Pol II levels, achieved with biallelic degron at high 5-Ph-IAA concentrations, did transcription begin to decline (Fig. S2D, Fig. S4D). Notably, the single-cell variability falls within this buffering window, suggesting it may enable transcriptional homeostasis despite cell-to-cell variation in Pol II levels. This robustness contrasts with a limiting factor model, where such variation would directly translate into transcriptional variability.

### The nucleoplasmic pool adjusts to changes in Pol II abundance

To determine how cells maintain stable transcription despite reduced Pol II abundance, we examined the distribution of polymerases between chromatin-bound and nucleoplasmic pools. Actively elongating Pol II stably binds chromatin and can be distinguished from freely diffusing or transiently bound Pol II in live cells using fluorescence recovery after photobleaching (FRAP) (*15, 20, 21*). We performed FRAP on mAC-POLR2A in biallelic degron cells (Fig. S5) after 4 hours of treatment with either low (10 nM) or high (1 µM) concentrations of 5-Ph-IAA, when Pol II abundance is relatively stable.

At low 5-Ph-IAA concentration, nuclear mClover fluorescence declined by 30 ± 2%, yet this resulted in a 56 ± 2% reduction in the free pool, while chromatin-bound POLR2A declined by only 14 ± 6%. This disproportionate reduction suggests cells preferentially preserve chromatin-bound Pol II by drawing from the nucleoplasmic pool. At high 5-Ph-IAA concentration, Pol II levels were reduced by 69 ± 3%, which resulted in almost complete depletion of the nucleoplasmic pool (Fig. 2A). Under these conditions, chromatin-bound POLR2A also declined substantially (56 ± 2%), consistent with the pronounced reduction in pSer2 and 5-EU incorporation (Fig. 1D, Fig. S2C-D). This indicates that once the nucleoplasmic pool is exhausted, chromatin-bound Pol II can no longer be maintained.

**Figure 2.**
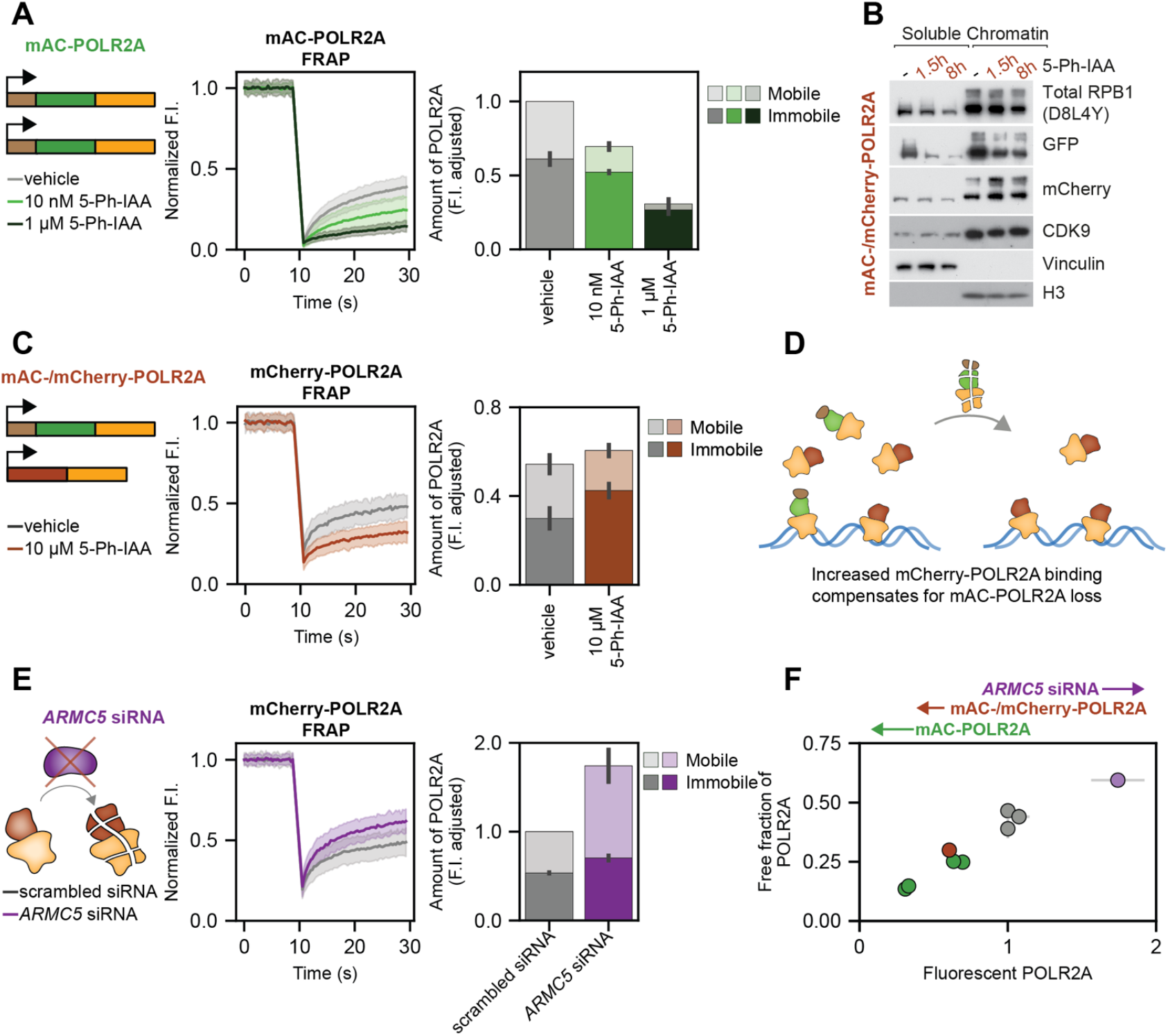
The nucleoplasmic pool adjusts to changes in Pol II abundance. (**A**) mAC-POLR2A FRAP in biallelic mAC-POLR2A cells and quantification of fluorescence-intensity-adjusted mobile and immobile fractions (45 cells collected across three experiments). (**B**) Chromatin fractionation and western blot. (**C**) As in A for mCherry-POLR2A FRAP in monoallelic mAC-/mCherry-POLR2A degron cells (45 cells collected across three experiments). (**D**) mCherry-POLR2A binding increases following mAC-POLR2A depletion. (**E**) As in A, for mCherry-POLR2A FRAP in ARMC5-depleted cells (30 cells collected across three experiments). (**F**) Mobile fraction (nucleoplasmic) from FRAP as a function of total Pol II abundance, measured as mean nuclear intensity of fluorescent POLR2A. Note some error bars do not extent beyond points and are not visible. All kinetic FRAP traces are shown as mean ± S.D. of all cells collected in three experiments. Quantification of Pol II fractions and summary plots are shown as mean ± S.D. of three separate experiments.

To confirm that this represents genuine redistribution rather than selective protection of chromatin-bound Pol II in mAC-POLR2A degron cells, we turned to monoallelic degron cells, where we can monitor the non-degraded mCherry-POLR2A pool while depleting mAC-POLR2A. Following 90 minutes of 5-Ph-IAA treatment, FRAP revealed a 44 ± 19% increase in chromatin-bound mCherry-POLR2A (Fig. 2C-D), which was confirmed by chromatin fractionation (Fig. 2B). Conversely, ARMC5 knockdown increased Pol II levels by 74 ± 18%, but FRAP measurements showed this excess accumulated predominantly in the nucleoplasmic pool, which increased by 123 ± 26% (Fig. 2E).

Across all perturbations, the fraction of Pol II which is nucleoplasmic and mobile scaled with total Pol II levels (Fig. 2F), demonstrating that the free pool serves as a dynamic buffer that contracts when Pol II is depleted to preserve chromatin-bound polymerases and expands when Pol II is elevated to absorb excess.

### The promoter-proximal checkpoint buffers changes in Pol II abundance

Having established that the nucleoplasmic pool buffers changes in Pol II abundance, we next asked whether chromatin-bound polymerases also redistribute across different stages of the transcription cycle. Before entering productive elongation, Pol II passes through the promoter-proximal region where it may pause or terminate rather than continuing (*6*), a major decision point where polymerase usage efficiency could be adjusted. Promoter-proximal Pol II is marked by Pol II CTD phosphorylation at Serine 5 (pSer5). To examine whether this population changes when Pol II is depleted, we quantified pSer5 levels in monoallelic POLR2A degron cells after 90 min of 5-Ph-IAA treatment. pSer5 decreased by 11 ± 4%, which was intermediate between total POLR2A (25 ± 3% decrease) and pSer2 (2 ± 5% decrease) (Fig. 3A), suggesting promoter-proximal Pol II is partially buffered.

**Figure 3.**
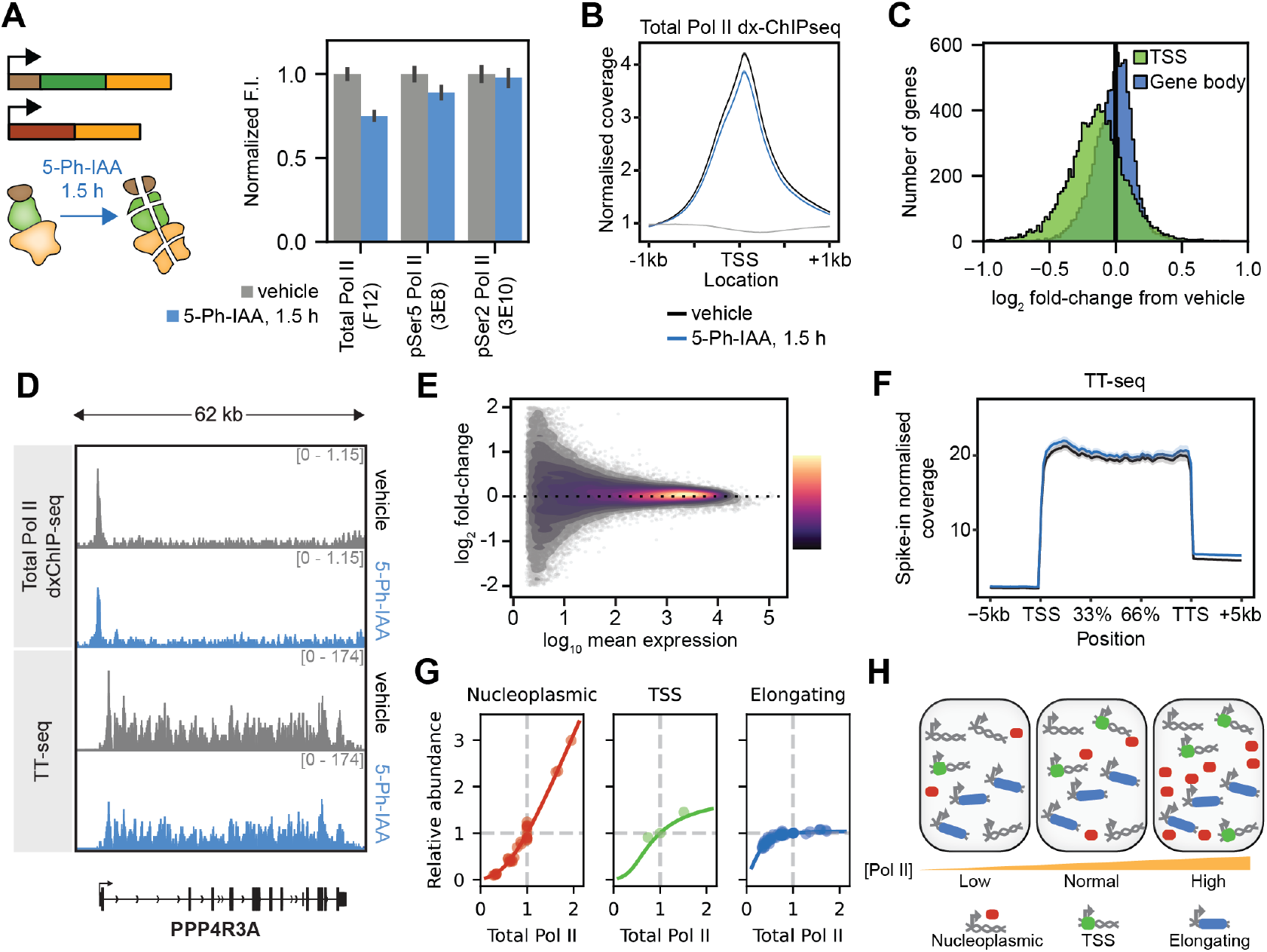
The promoter-proximal checkpoint buffers changes in Pol II abundance. (**A**) Total Pol II (F12), pSer5 Pol II (3E8) and pSer2 Pol II (3E10) immunofluorescence in mAC-/mCherry-POLR2A cells. Shown as mean ± S.D. of replicate wells collected in three separate experiments. (**B**) dxChIP-seq metagene profiles of RNA Pol II occupancy at the TSS. (**C**) log_2_ fold-change in normalized coverage in total Pol II dxChIP-seq at the TSS and gene body. (**D**) Example gene track for total Pol II dxChIP-seq and TT_chem_-seq. (**E**) Spike-in normalized TT_chem_-seq results for individual genes. Points colored by point density. (**F**) Spike-in normalized TT_chem_-seq metagene profile. Legend as in B. (**G**) Mathematical model of Pol II pools together with experimental data from FRAP mobile amount, (Nucleoplasmic), dxChIP-seq (TSS), pSer2 immunofluorescence (Elongating). (**H**) As total Pol II levels are manipulated, amount of Pol II in the nucleoplasmic pool and bound in the promoter-proximal region buffers elongating Pol II. All panels show comparison of vehicle treatment to 1.5 hours of 10 µM 5-Ph-IAA treatment in monoallelic mAC-/mCherry-POLR2A degron cells.

To examine Pol II occupancy genome-wide, we performed dxChIP-seq for total POLR2A in monoallelic degron cells after 90 min of 5-Ph-IAA treatment. This revealed a global decrease in Pol II at promoter-proximal regions, but not in gene bodies or transcription termination sites (Fig. 3B-D, Fig. S6A-B). This indicates that polymerase redistribution occurs not only between nucleoplasmic and chromatin-bound pools, but also between promoter-proximal and elongating states. To examine transcriptional output, we performed spike-in normalized transient-transcriptome sequencing (TT_chem_-seq) (*22*), which quantifies nascent RNA genome-wide. TT_chem_-seq revealed no significant changes in nascent RNA levels for individual genes despite reduced total Pol II (Fig. 3E, Methods). Metagene plots confirmed uniform transcriptional output from transcription start sites through to termination regions (Fig. 3F). Thus, the same number of polymerases progress to elongation despite reduced promoter-proximal occupancy, consistent with stable pSer2 Pol II, EU incorporation, and gene-body POLR2A levels. Notably, we previously observed the opposite pattern when Pol II abundance *increases* due to ARMC5 knockout (*15*). In that case, Pol II preferentially accumulates in the promoter-proximal zone while gene-body Pol II levels and nascent RNA remain relatively unchanged. Together, these observations demonstrate bidirectional buffering: the promoter-proximal pool shrinks when Pol II is reduced and expands when Pol II is increased, while in both cases transcriptional output remains constant.

How do cells maintain constant transcriptional output from a reduced promoter-proximal pool? Two broad classes of mechanisms could account for this. In an ‘active’ model, cells would sense reduced Pol II levels and compensate by modulating transition rates – for example, increasing P-TEFb activity to increase the probability of Pol II transitioning to elongation and thereby maintaining constant elongation flux from a diminished promoter-proximal pool. In a ‘passive’ model, transition rates in the transcription cycle may be limited by factors other than Pol II availability, so that flux through these transitions would not linearly depend on Pol II pool size and no sensing or adaptive rate modulation would be required. Constant nascent RNA (TT_chem_-seq) and gene-body Pol II (dxChIP-seq) indicate that the number of Pol II released per unit time remains unchanged. Therefore, reduced promoter-proximal Pol II means either that the rate of polymerase entry decreases, or the rate of premature termination increases.

To test whether a passive model can account for our observations, we developed a mathematical model of the transcription cycle with sequential transitions governing chromatin-binding, premature termination, release into elongation, and termination (Methods). We assigned Michaelis-Menten kinetics to chromatin-binding and release into elongation. Fitting this model to the full range of Pol II perturbations – including biallelic and monoallelic POLR2A depletion and ARMC5 disruption – using measurements of nucleoplasmic Pol II from FRAP, promoter-proximal Pol II from dxChIP-seq, and pSer2 levels from immunofluorescence, revealed that low Michaelis constants (K_m_) could recapitulate these observations (Fig. 3G). With low K_m_, these steps operate near saturation (analogous to enzymes working at maximum capacity) so that reduction in Pol II does not proportionally reduce output. Such passive buffering requires no active sensing or rate modulation.

We next tested whether active modulation of the promoter-proximal checkpoint contributes to buffering by examining P-TEFb, the kinase responsible for Ser2 phosphorylation and release into productive elongation (*8*). P-TEFb is sequestered by the 7SK ribonucleoprotein (RNP) complex, which has been hypothesized to globally regulate release into elongation (*23*). We observed no global changes in P-TEFb subunit abundance or chromatin binding following partial Pol II depletion (Fig. 2B, Fig. S6C). Knockdown of 7SK RNP components HEXIM1/2 or LARP7 slightly increased baseline POLR2A and pSer2 levels but did not affect stability of pSer2 levels during mAC-POLR2A depletion in monoallelic degron cells (Fig. S7, S8). This indicates that neither 7SK-mediated P-TEFb regulation, nor P-TEFb abundance and chromatin binding, contribute to maintaining elongating Pol II when Pol II abundance decreases.

While we cannot rule out other adaptive mechanisms such as modulation of premature termination or other early elongation factors, a passive kinetic buffering model is consistent with all available data. Together, these results indicate that Pol II activity is buffered bidirectionally by both chromatin binding and release to elongation, maintaining constant transcriptional output across a wide range of abundance changes (Fig. 3H). This robustness can be achieved through saturated kinetics without requiring active sensing of Pol II levels or modulation of kinetic rates, indicating that the multi-checkpoint structure of the transcription cycle provides inherent buffering capacity.

### Pol II levels recover following monoallelic POLR2A depletion

Having observed stable transcription within 1.5 hours of partial Pol II depletion, we next examined longer timescales in monoallelic degron cells. Surprisingly, beyond 2 hours of mAC-POLR2A degradation, we observed a substantial increase in mCherry-POLR2A levels (35 ± 7% at 24 h, Fig. 4A), which progressively restored total Pol II levels to 89 ± 10% of baseline within 8 hours. FRAP measurements after 24 hours showed that absolute bound amounts of mCherry-POLR2A, expressed from a single allele, matched levels in cells with biallelic mCherry-POLR2A (Fig. 4B, Fig. S9A), and that nucleoplasmic Pol II was largely restored (Fig. 4C).

**Figure 4.**
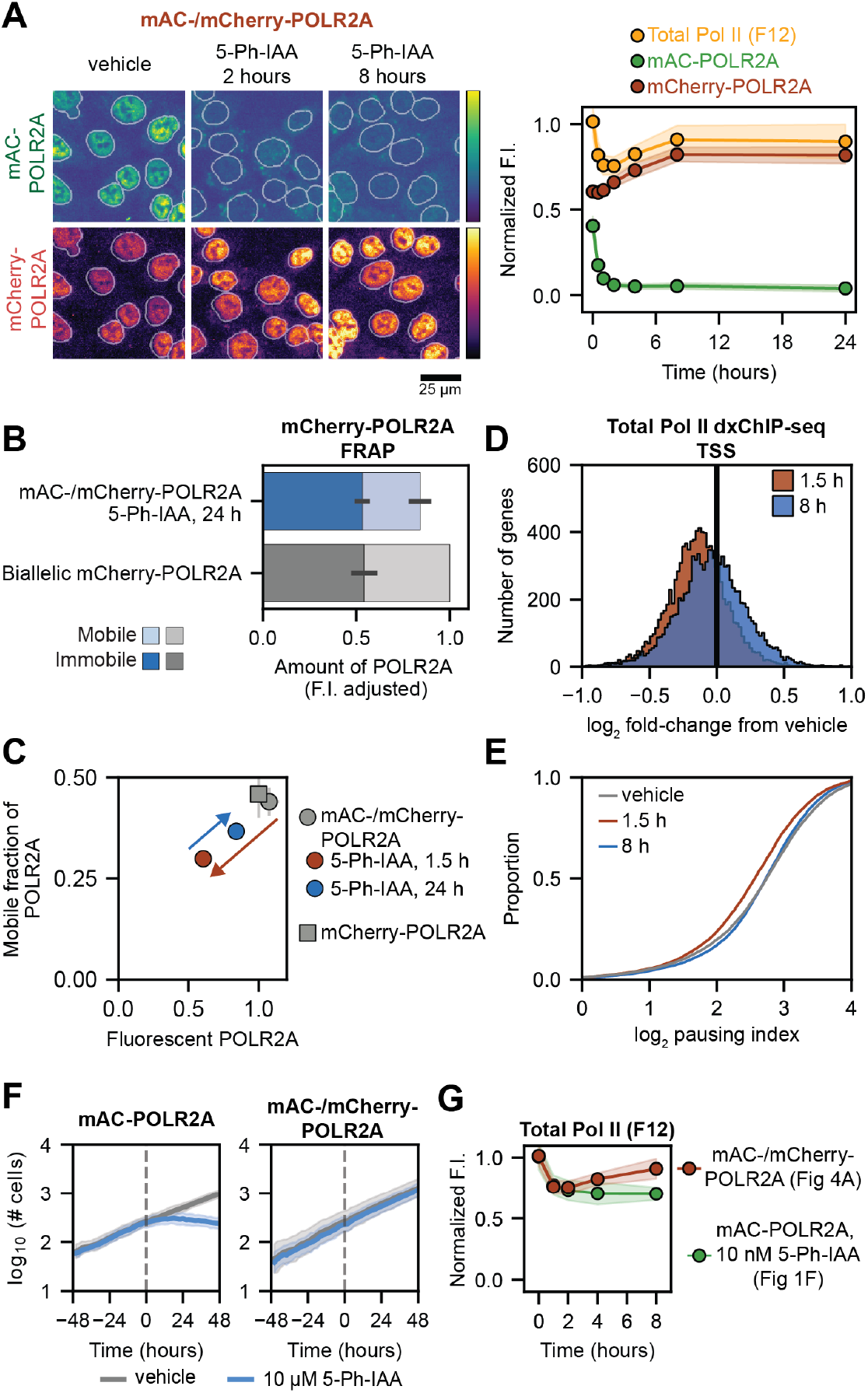
Pol II levels recover following monoallelic POLR2A depletion. (**A**) mClover, mCherry fluorescence and total Pol II immunofluorescence in monoallelic mAC-/mCherry-POLR2A degron cells. Example images and quantification of time course after 5-Ph-IAA addition, normalised to fluorescence levels in biallelic mAC-POLR2A cells and biallelic mCherry-POLR2A cells, respectively. Shown as mean ± S.D. of replicate wells collected in three separate experiments. (**B**) Amount of bound and free POLR2A in mAC-/mCherry-POLR2A cells treated with 5-Ph-IAA for 24 hours, compared to biallelic mCherry-POLR2A cells. Shown as mean ± S.D. of three separate experiments. (**C**) Total POLR2A levels and mobile fraction of POLR2A in mAC-/mCherry-POLR2A cells treated with 5-Ph-IAA at different timepoints, compared to biallelic mCherry-POLR2A cells. (**D**) dxChIP-seq of log_2_ fold-change in total Pol II TSS coverage after 1.5 or 8 hours of 5-Ph-IAA. (**E**) Empirical cumulative distribution function of pausing index of all genes with pausing index greater than 2 and TSS coverage greater than 0.5 after vehicle, or either 1.5 hours or 8 hours 5-Ph-IAA treatment. (**F**) Growth rates of biallelic mAC-POLR2A and mAC-/mCherry-POLR2A cells, treated with either vehicle or 5-Ph-IAA at indicated grey line. Shown as mean ± S.D. of replicate wells collected in two separate experiments. (**G**) Comparison of total Pol II abundance via immunofluorescence in mAC-POLR2A cells following treatment with low 5-Ph-IAA concentration, and mAC-/mCherry-POLR2A cells following high 5-Ph-IAA concentration, reproduced from earlier figures.

To test whether this recovery also restores the normal Pol II distribution across transcription cycle stages, we performed Pol II dxChIP-seq after 8 hours of 5-Ph-IAA treatment, an earlier timepoint where recovery was mostly complete. The global shift to lower promoter-proximal Pol II seen at 1.5 hours was reversed, with average TSS Pol II occupancy returning to near-baseline while gene-body levels remained unchanged (Fig. 4D, Fig. S9B). Consistent with this, the pausing index (*24, 25*), which measures relative Pol II occupancy at promoters versus gene bodies, decreased at 1.5 hours when total Pol II was lowest, then reverted to baseline by 8 hours (Fig. 4E). This restoration of nucleoplasmic and promoter-proximal Pol II suggests that recovery on hour timescales provides a second layer of robustness, complementing the immediate transcriptional buffering occurring within minutes. Growth rates remained unaltered during monoallelic POLR2A depletion (Fig. 4F, Fig. S9C), indicating that these combined mechanisms fully buffer the perturbation.

Notably, in biallelic degron cells depleted to similar Pol II levels, no recovery of nuclear POLR2A abundance occurred (Fig. 4G), revealing that the compensation mechanism depends on having one intact POLR2A allele available. What mechanisms might account for the increase in mCherry-POLR2A upon mAC-POLR2A depletion? Increased nuclear POLR2A protein could arise through changes at the RNA level (increased mRNA synthesis or decreased mRNA degradation) or at the protein level (increased translation, assembly, or nuclear import rates, or decreased protein degradation at any of these stages) (Fig. 5A). To distinguish these possibilities, we first measured *POLR2A* mRNA levels using single-molecule RNA fluorescence in situ hybridization (smFISH) in monoallelic mAC/mCherry-POLR2A cells. 5-Ph-IAA treatment for 24 hours did not alter transcript abundance (Fig. 5B-C, Fig. S9D-F). We next measured nuclear mCherry-POLR2A protein stability in live cells using fluorescence bleach-chase (*26*), which revealed no change in half-life following 5-Ph-IAA treatment (Fig. 5D, Fig. S10A). Consistent with recovery arising through cytoplasmic protein synthesis, assembly, or import, inhibiting protein translation with cycloheximide prevented mCherry-POLR2A accumulation in response to mAC-POLR2A loss (Fig. 5E). Inhibiting POLR2A nuclear import with leptomycin B (*27*) also prevented mCherry-POLR2A accumulation (Fig. S12A). Together, these results rule out changes in transcription of *POLR2A*, mRNA stability, and nuclear protein stability, indicating that recovery operates through cytoplasmic processes: translation, assembly, or import.

**Figure 5.**
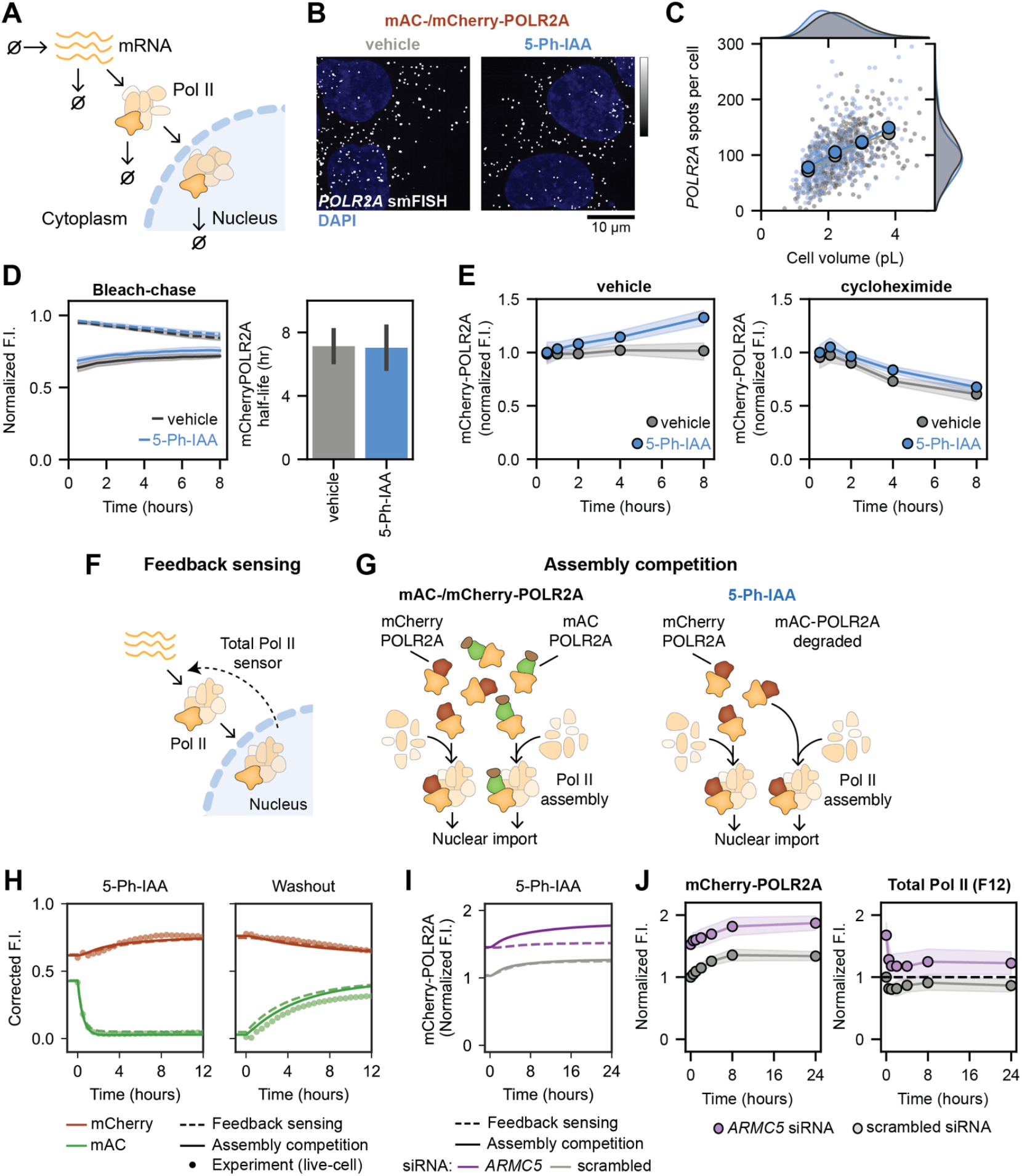
Excess subunit production confers robustness of Pol II levels. (**A**) Schematic of major steps in Pol II biosynthesis. (**B**) Example images of smFISH in mAC-/mCherry-POLR2A degron cells after 24 h of 5-Ph-IAA treatment. (**C**) Spot count as a function of cell volume. (**D**) Bleach-chase assay to measure mCherry-POLR2A half-life upon mAC-POLR2A depletion. Shown as mean ± S.D. of replicate wells collected in three separate experiments. (**E**) Cycloheximide addition prevents 5-Ph-IAA-induced mCherry-POLR2A increase in monoallelic degron cells. (**F**) Schematic of feedback sensing mechanism, where Pol II biosynthesis depends on nuclear Pol II levels. (**G**) Schematic of assembly competition mechanism. POLR2A subunits compete for incorporation into nuclear Pol II holoenzyme. Competition is relieved upon loss of mAC-POLR2A. (**H**) Live-cell imaging of mCherry and mClover during 5-Ph-IAA treatment (left) and washout (right) in monoallelic degron cells, together with model fits. (**I**) Model predictions from altered nuclear Pol II half-life, through ARMC5 depletion. (**J**) Increase of mCherry-POLR2A and recovery of total Pol II immunofluorescence intensity during 5-Ph-IAA treatment in ARMC5-depleted cells.

These constraints are consistent with two distinct classes of mechanisms. In a regulated ‘feedback sensing’ model (Fig. 5F), cells would detect reduced POLR2A levels (for example, by monitoring nucleoplasmic Pol II concentration) and increase POLR2A translation or nuclear import. Alternatively, in a constitutive ‘assembly competition’ model (Fig. 5G), excess production of POLR2A subunits would compete for a limiting factor required for Pol II holoenzyme assembly or nuclear import. Here, mAC-POLR2A depletion would shift the competitive balance toward increased mCherry-POLR2A incorporation and import without any sensing mechanism.

To determine whether these models could reproduce our observations, we formulated them mathematically and performed computational simulations (Methods). For assembly competition, we modelled constitutive POLR2A production, a limiting assembly/import factor, and cytoplasmic degradation of unassembled subunits. For feedback sensing, we implemented proportional feedback where Pol II synthesis increases as nuclear Pol II levels drop. Both models reproduced mCherry-POLR2A and mAC-POLR2A dynamics observed in live cells during 5-Ph-IAA treatment and washout (Fig. 5H). Washout experiments showed mCherry-POLR2A declining before total Pol II fully recovered, inconsistent with set-point control that would maintain elevated mCherry-POLR2A until normal Pol II levels are restored. To distinguish between feedback sensing and assembly competition, we tested a condition where total Pol II levels would remain elevated during mAC-POLR2A depletion.

We depleted ARMC5 to elevate Pol II above normal levels, then induced mAC-POLR2A degradation. The feedback sensing model predicts no mCherry-POLR2A increase when total Pol II exceeds normal levels because ARMC5 knockdown was previously shown to increase POLR2A stability without affecting synthesis (*15*). In contrast, assembly competition predicts mCherry-POLR2A increase whenever mAC-POLR2A becomes depleted relative to mCherry-POLR2A, independent of total Pol II levels (Fig. 5I). mCherry-POLR2A increased similarly in ARMC5-depleted cells as in controls (22 ± 6% versus 34 ± 7% at 24 h), despite total Pol II remaining above normal levels throughout (Fig. 5J), supporting assembly competition over feedback sensing. This mechanism also explains why biallelic degron cells fail to recover (Fig. 4G) and is consistent with observations that POLR2A is less stable early in its lifetime and stabilizes with age (*28*), the expected behavior for subunits produced in excess and degraded unless incorporated into stable complexes.

Together, these results indicate that cells maintain Pol II abundance homeostasis through constitutive excess subunit production and assembly competition rather than feedback sensing. Operating on hour timescales, this mechanism restores both total Pol II levels and the normal distribution between nucleoplasmic and chromatin-bound pools, providing a second layer of buffering beyond the immediate transcriptional compensation that occurs within minutes.

### Sustained Pol II depletion reveals additional compensatory mechanisms

The buffering mechanisms identified above operate within hours and maintain Pol II levels and transcriptional output across modest perturbations. Under more severe and sustained Pol II depletion (biallelic degron, 10 µM 5-Ph-IAA for 4-24 h), which decreased transcription and abolished cell growth (Fig. 4F, Fig. S2, S11A), additional compensatory responses emerged that were not observed with milder perturbations.

*POLR2A* transcript levels increased substantially after 4 h of POLR2A depletion (140 ± 20% increase at 24 h), while *POLR2B* and *POLR2C* transcripts remained unchanged (Fig. S11). This selective increase did not occur with general transcriptional inhibitors but was observed when inhibiting Pol II assembly with CB6644, which targets the R2TP chaperone complex (*29*) (Fig. S12). Additionally, P-TEFb subunits CDK9 and CCNT1 were progressively upregulated (35 ± 12 % increase at 4 h, 84 ± 22% increase at 24 h for CDK9), a response also observed in independent POLR2B knockdown experiments (Fig. S13, S7A).

These late-acting responses reveal additional layers of compensation that activate under severe perturbations, suggesting a hierarchical buffering system capable of responding to escalating stress.

## Discussion

Robustness to fluctuations in essential machinery can emerge from the architecture of biosynthetic pathways themselves, without requiring dedicated sensing mechanisms. This is observed in metabolic pathways where flux through saturated enzymes becomes independent of substrate availability (*30*). Our results reveal that human cells maintain transcriptional homeostasis through hierarchical buffering mechanisms in which saturated kinetics at multiple stages confer stability. Within minutes, chromatin-binding and pause release checkpoints operating near saturation enable constant transcriptional output despite changes in Pol II availability. Over hours, constitutive excess POLR2A subunit production creates competition for incorporation into Pol II holoenzyme, restoring total Pol II levels without feedback sensing. This assembly competition mechanism may be a widespread buffering strategy, as rapid degradation of unincorporated subunits is common across essential multiprotein complexes (*28, 31*).

Our findings contrast fundamentally with yeast, where Pol II appears limiting and Pol II chromatin-binding scales directly with Pol II abundance (*5*). Human cells instead maintain substantial Pol II excess distributed across nucleoplasmic and promoter-proximal pools. The evolution of additional checkpoints in metazoan transcription, such as promoter-proximal pausing (*32*), may have enabled this shift. These create additional buffers that can expand or contract without altering transcriptional output. This strategy trades the metabolic cost of excess polymerase production for increased robustness to perturbations, which may be critical for maintaining transcriptional precision across diverse cell types in multicellular organisms.

Why cells maintain such buffers becomes clearer in light of recent work identifying nucleoplasmic Pol II depletion as an apoptotic trigger independent of transcriptional inhibition (*12*). Our observation that cells prioritize preserving chromatin-bound polymerases through multiple mechanisms is consistent with the nucleoplasmic pool functioning not only as a quantitative buffer but also as a sensor of biosynthetic capacity. When perturbations exhaust this reservoir, cells may interpret the signal as catastrophic proteostatic failure rather than simply reduced transcription. This has implications for understanding how transcriptional inhibitors and other chemotherapies, which also use this mechanism, preferentially target cancer cells (*28, 31*)

## Acknowledgements

S.B. is the recipient of an Australian Research Council Discovery Early Career Award (DE230100271) funded by the Australian government. A.T.V. is supported by a core grant to the LMB from the Medical Research Council (MC_UP_1201/28). We thank UNSW’s Katharina Gaus Light Microscopy Facility, Flow Cytometry, and Research Technology Services as well as MRC LMB Mechanical and Electronics Workshops and Scientific Computing for supporting our work.

## Materials and Methods

### Cell lines and growth conditions

HCT116 cells (male) were obtained from ATCC (CCL-247). HCT116 OsTIR(F74G) express OsTIR(F74G) from the safe-harbor AAVS1 locus from a constitutive CMV promoter (*13, 14*). These were a gift from Masato Kanemaki. Puromycin selection marker was excised by Cre-mediated recombination. Cells were maintained in McCoy’s 5A modified medium (Gibco 16600108) in 10% FBS (Moregate Biotech) at 37 ºC and 5% CO2.

### Generation of knock-in cell lines

HCT116 cells with biallelic mCherry-POLR2A were described previously (*15*). HCT116 OsTIR(F74G) with biallelic mAID-mClover-POLR2A, monoallelic mAID-mClover-POLR2A / mCherry-POLR2A and biallelic ARMC5-mAID-mClover-Hyg were generated using CRISPR-mediated homology-directed repair. The N-terminal mAID-mClover-POLR2A donor plasmid was generated by removing the Hygromycin resistance gene from mAC-POLR2A donor (Addgene #124496) using Gibson assembly. The C-terminal ARMC5-mAID-mClover-Hyg donor was generated by Gibson assembly, amplifying fragments by PCR from HCT116 genomic DNA, and pMK290 (mAID-mClover-Hygro (*33*) ; Addgene #124496). The assembled ARMC5 donor plasmid contains a 614bp left homology arm (not including the stop codon), followed by a Ser-Gly-Ala-Gly-Ala linker then the mini-AID tag, mClover3 protein, SV40 terminator and then PGK promoter driving HygR, and finally a 562 bp right homology arm. The pX330 Cas9/gRNA plasmid previously used to generate mCherry-POLR2A cells (*13*) (Addgene #124495) was again used for all POLR2A knock-in cells. C-terminal ARMC5-specific gRNA sequences were cloned into pX459 Cas9 plasmid (*34*) and used to generate biallelic ARMC5-mAID-mClover-Hyg cells. 7-10 days after transfection of plasmids, GFP-positive or GFP- and RFP-positive cells were selected via FACS and clonal populations grown from single cells. All knock-in cell lines were confirmed by PCR-based genotyping and Sanger sequencing.

### Cell growth assays

Parental HCT116 OsTIRF74G, mAC-/mCherry-POLR2A, or mAC-POLR2A cells were seeded at an equal density of 3000 cells per well in a clear plastic 96 well plate (Corning 3516) in McCoy’s 5A medium without phenol red (Cytiva SH30270.01) supplemented with 10% FBS (Moregate Biotech) and 1% penicillin/streptomycin (Sigma Aldrich P0781). Shortly after plating, bright-field imaging was begun with a 20x air objective on a Sartorius IncuCyte SX5, contained within a tissue-culture incubator at 37 ºC with 5% CO_2_. Four fields of view were collected for each well every 2 hours for 48 hours, at which point either 5-Ph-IAA or vehicle was added at a 10x concentration, 22 µL onto 200 µL, for a final of 10 µM 5-Ph-IAA in 0.1% DMSO in imaging media, or vehicle only. Imaging continued for a further 48 hours after compound addition.

Image processing was performed using custom pipeline written in python. Images were segmented using Cellpose3 (*35*) with the built-in model ‘livecell_cp3’. Total number of cells were summed over fields of view for a total count per well over time. Two separate experiments were performed, with 4 - 5 replicates each of vehicle and 5-Ph-IAA treated wells collected in each experiment. log_10_ values cell count over time are shown as mean ± S.D of replicate wells. Growth rates were determined per-well by simple linear regression using the scipy package to the log_10_ of cell count over time and grouped by treatment condition.

### Preparation of cells for microscopy

For high-throughput imaging, cells were plated in microscopy-compatible clear plastic 96-well plates (Greiner µClear 781091, bleach-chase and live imaging of fluorescent proteins) or 384-well plates (Greiner µClear 781091, immunofluorescence, 5-EU, or 7SK FISH assays). For smFISH assays, 18-well glass-bottom plates (ibidi 81817) were incubated with fibronectin (Merck Sigma Aldrich 341631) at 20 µg/mL for over 3 hours at 37 ºC, with fibronectin solution aspirated immediately before cell addition. For FRAP assays, 8-well glass-bottom ibidi plates (ibidi 80827) were incubated with 100 µg/mL of poly-L-lysine for 1 hour at room temperature, rinsed 3x with PBS, and allowed to air dry in a tissue culture hood before cell addition. In all cases cells were allowed to settle in plates before returning to incubator.

For all fixed cell imaging (immunofluorescence, 5-EU, 7SK FISH, and smFISH assays) cells were seeded in McCoy’s 5A modified medium (Thermo Fisher Gibco 16600108) supplemented with 10% FBS (Moregate Biotech) (‘media’). For live-cell imaging (bleach-chase, live imaging of fluorescent proteins, and FRAP assays), cells were seeded in McCoy’s 5A without phenol red (Cytiva SH30270.01) supplemented with 10% FBS (Moregate) (‘imaging media’).

For high-throughput fixed-cell imaging assays, 4x washes in either PBS (immunofluorescence and 5-EU assays) or 2x SSC (7SK FISH assay) were performed between each step, and additions occurred by adding 30 µL of 2x solutions onto a 30 µL residual volume, using a plate wash/dispenser (Biotek EL406). For immunofluorescence, 5-EU, and 7SK FISH assays, cells were fixed in 4% paraformaldehyde freshly diluted from ampoules (EMS Emgrid 15710) for 15 minutes. For smFISH assay and fixed-cell FRAP control imaging, media was completely aspirated from 8-well or 18-well plates before addition of 4% freshly diluted paraformaldehyde for 15 minutes.

For immunofluorescence and 5-EU assays, cells were permeabilized in 0.25% Triton X-100 (Sigma Aldrich 93443) in PBS for 10 minutes. For 7SK FISH and smFISH assays, cells were permeabilised in 70% ethanol at 4 ºC for 4-6 hours.

### Live-cell imaging of fluorescent proteins

Cells were plated at a density of 4000 cells per well in imaging media. 8.5 hours before imaging commenced, imaging media was exchanged for fresh imaging media additionally containing 1% penicillin/streptomycin (Sigma Aldrich P0781), and either 5-Ph-IAA or vehicle was added. 0.5 hours before imaging commenced, imaging media was completely aspirated and replaced 4x, before a final addition of 200 µL of imaging media, followed by either vehicle or 5-Ph-IAA at 10x concentrations. Imaging was then performed on a Nikon Ti2 microscope with Yokogawa CSU-W1 spinning disk using a 20x, 0.75NA objective and Hamamatsu ORCA-Fusion C14440-20UP cameras, equipped with a Tokai Hit heated stage and chamber, at 37 ºC with 5% CO_2_, and humidity controlled. 7x z-planes were acquired at 2 µm intervals of miniAID-mClover-POLR2A, 488 nm laser and 525/50 filter, and mCherry-POLR2A, 561 nm laser and 617/73 filter, sequentially at 15-minute intervals. 9 imaging sites were collected across four separate conditions – initially treated with vehicle or 5-Ph-IAA, washed and followed by either vehicle or 5-Ph-IAA – in single wells in two experiments.

Image processing was performed on maximum projected images using a custom pipeline written in python. Nuclei were segmented on mCherry-POLR2A signal using a using a manually trained Cellpose 2.0 model, based on the built-in ‘nuclei’ model. Mean mCherry-POLR2A and mAC-POLR2A fluorescence of each nucleus was taken and averaged per replicate well. Acquisition photobleaching over the time-course of the experiment was apparent to the extent of ∼25%, and was corrected by fitting to an exponential decay function using scipy. mCherry-POLR2A intensity was background subtracted using an estimate from a region outside the cells and normalized to the initial timepoint of a vehicle-only well within that experiment as 1. mAC-POLR2A intensity was background subtracted using 5-Ph-IAA treated wells and normalized to the initial timepoint of a vehicle-only well within that experiment as 1.

### Bleach-chase experiments (protein half-life measurement)

Bleach-chase experiments were performed essentially as previously described (*15*). Cells were plated at a density of 4000 cells per well in imaging media. 8 hours before imaging commenced, imaging media was exchanged for fresh, additionally containing 1% penicillin/streptomycin (Sigma Aldrich P0781), and 5-Ph-IAA or vehicle was added. Imaging was then performed on a Nikon Ti2 microscope with Yokogawa CSU-W1 spinning disk using a 20x, 0.75NA objective and Hamamatsu ORCA-Fusion C14440-20UP camera, equipped with a Tokai Hit heated stage and chamber, at 37 ºC with 5% CO_2_, and humidity controlled. 7x z-planes were acquired of mCherry-POLR2A at 2 µm steps at a 30-minute interval using a 561 nm laser and 617/73 nm filter, collecting both control regions or bleach regions within each well, 9 imaging sites at each. After 8 loops (4 hours) of imaging all wells, bleach steps were performed over 7x z-planes in 2 µM steps using a widefield light source from a mercury vapor lamp with 635/60 nm filter, for 4 loops. Following bleaching, both bleached and control regions of each well were imaged for a further 16 loops (8 hours). Two replicate wells of both vehicle- and 5-Ph-IAA-treated cells were imaged in each of three separate experiments.

Image processing was performed on maximum projected images using custom pipeline written in python. Nuclei were segmented on mCherry-POLR2A signal using a using a manually trained Cellpose 2.0 model, based on the built-in ‘nuclei’ model. Mean mCherry-POLR2A fluorescence of each nucleus was taken, and background fluorescence intensity estimated from a region outside the cells and subtracted from intensity values. mCherry-POLR2A fluorescence intensity was then normalised to the pre-bleach step baseline per bleached or control region, per well. Log-transformed mean fluorescence intensity of the bleach region was subtracted from that of the control region within each well to estimate the decay of the invisible, bleached mCherry-POLR2A (Fig. S10A). These values were then fit to a linear least-squares regression using the scipy package to estimate total mCherry-POLR2A turnover. Growth rates were calculated per bleached or control region, per well, by taking log-transformed values of the number of nuclei counted across all fields of view through time and performing a linear least-squares regression using the scipy package. No effect of bleaching on cell growth was observed. Dilution of mCherry-POLR2A due to cell growth was subtracted from total mCherry-POLR2A removal rate to estimate degradation rate for either vehicle or 5-Ph-IAA treated cells.

### Immunofluorescence

After fixation and permeabilization, cells were blocked in 50% blocking buffer (Millenium Biosciences Li-Cor Intercept in PBS, LCR-927-70001) in PBS for 30 minutes. Wells were incubated with primary antibodies in 50% blocking buffer for 90 minutes. Secondary antibodies then incubated for 30 minutes in 50% blocking buffer. Where necessary, DAPI at 100 ng/mL final was added together with secondary antibodies. Where multiple stains were performed, primary and secondary antibodies were added in series.

### 5-ethynyl uridine visualization via click chemistry

After fixation and permeabilization, cells were washed into Tris-buffered saline (125 mM sodium chloride, 50 mM Tris pH 8 Thermo Fisher Invitrogen AM9856). A 1.5x click reaction mixture was made in TBS containing 150 mM sodium ascorbate (Sigma Aldrich A7631), 3 mM copper sulphate (Chem-Supply Australia CA068) and 7.5 mM Alexa647 azide (Thermo Fisher Invitrogen A10277) or AF647 azide (Lumiprobe A6830). 30 µL of 1.5x click reaction was added onto 15 µL residual TBS and incubated for 30 minutes at room temperature. Where necessary, DAPI at final 100 ng/mL was added for 5 minutes following click reaction.

### 7SK fluorescence in-situ hybridization (FISH)

After fixation and permeabilization, cells were exchanged into a wash buffer of 2x SSC (Thermo Fisher Invitrogen AM9763) and 10% formamide (Thermo Fisher Invitrogen AM9342) via a 2x 10-minute wash, 90 µL onto 15 µL residual. Previously published oligonucleotide 7SK FISH probes (*36*) labelled with ATTO647N (IDT) were then added 45 µL onto 15 µL residual for a 100 nM final concentration in a hybridization buffer of 2x SSC, 10% formamide, 2 mM ribonucleoside vanadyl complexes (New England Biolabs S1402S), 200 µg/mL BSA (Thermo Fisher Invitrogen AM2616), and 100 mg/mL dextran sulphate (Merck Sigma Aldrich D8906-50G). Cells were incubated with probes overnight in a humidity-controlled tissue culture incubator at 37 ºC. The next day, two one-hour washes were performed in 2x SSC, 10% formamide wash buffer, the second wash containing 100 ng/mL DAPI, at 37 ºC in the same incubator. After a final 10-minute, room temperature wash in the same wash buffer, a final wash into 2x SSC was performed, which the cells remained in for imaging.

### Single molecule fluorescence in-situ hybridisation (smFISH)

smFISH was performed essentially as described previously (*37*). smFISH probes were designed targeting *POLR2A, POLR2B*, and *POLR2C* using Stellaris FISH probe designer. Amine reactive ATTO565 N-hydroxysuccimide (Sigma Aldrich 72464-1MG) or ATTO633 N-hydroxysuccimide (Sigma Aldrich 1464-1MG) was reacted with amino-11-ddUTP (Lumiprobe 15040). 32 oligonucleotide probes (IDT) per set were pooled and labelled with either ATTO565-11-ddUTP or ATTO633-11-ddUTP using a terminal deoxytransferase (Thermo Fisher EP0162). Probes were then ethanol precipitated, purified using reverse-phase high-performance liquid chromatography to collect only fluorophore-labelled oligonucleotides, dried and isolated using an oligonucleotide clean-up kit (Zymo research R2052). For odd/even experiments, alternating probes from 64 total targeting *POLR2B* were pooled to two separate sets targeting the same transcript.

All wash and addition steps were performed with complete liquid aspiration from wells. After fixation and permeabilization, cells were exchanged into a wash buffer of 2x SSC (Thermo Fisher Invitrogen AM9763) and 10% formamide (Thermo Fisher Invitrogen AM9342) via a 2x 10-minute wash. Oligonucleotide smFISH probes were then added at 10 nM in a hybridisation buffer of 2x SSC, 10% formamide, 2 mM ribonucleoside vanadyl complexes, 200 µg/mL BSA, 100 mg/mL dextran sulphate, and 100 µg/mL yeast transfer RNAs (Thermo Fisher Invitrogen 15401011). Cells were incubated with probes overnight in a humidity-controlled tissue culture incubator at 37 ºC. The next day, two one-hour washes were performed in 2x SSC, 10% formamide wash buffer, the second wash containing 100 ng/mL DAPI, at 37 ºC in the same incubator. After a final 10-minute, room temperature wash in the same wash buffer, a 3x wash into 2x SSC was performed, which the cells remained in for imaging.

### siRNA transfection

For immunofluorescence and 7SK FISH experiments in 384-well plates, siRNAs were freshly diluted to a constant total siRNA concentration of 60 nM in Opti-MEM (Thermo Fisher Gibco 31985-062) and added 10 µL per well, followed by 10 µL of Lipofectamine RNAiMAX (Thermo Fisher Lipofectamine RNAiMAX 13778100) freshly diluted 1 in 125 in Opti-MEM. Transfection complexes were incubated at room temperature for 20 – 30 minutes, before cells were added in 50 µL of media. For most experiments, HCT116 cells were seeded at 1500 per well. For experiments where mAC-/mCherry-POLR2A cells were transfected with siRNAs, cells were plated at 2500 per well. Fixation was performed 3 days after plating, with 5-Ph-IAA time-course experiments offset such that time after siRNA transfection was maintained.

For single siRNAs, a final concentration of 30 nM of targeted siRNA and 30 nM of scrambled siRNA was used. For pooled siRNAs targeting a single gene, a final concentration of 10 nM of three individual siRNAs together with 30 nM of scrambled siRNA was used. For combinations of two sets of pooled siRNAs targeting two separate genes, a final concentration of 10 nM of six individual siRNAs, three for each gene, was used.

For RT-qPCR experiments in 6-well plates, 150 µL of pooled siRNAs at 30 nM total siRNA concentration, 10 nM of each individual, was added per well, followed by 150 µL of Lipofectamine RNAiMAX diluted 1 in 125. After 20 – 30 minutes room temperature incubation, HCT116 cells were seeded at 500 000 cells per well in 5 mL of media.

For FRAP experiments in 8-well plates, siRNAs were added 40 µL per well at 60 nM, followed by 40 µL of Lipofectamine RNAiMAX diluted 1 in 125. HCT116 mCherry-POLR2A cells were then added in 200 µL of imaging media at 8000 cells per well.

### Compound treatment

5-Ph-IAA (MedChemExpress HY-134653) was dissolved in DMSO at 10 mM and stored in aliquots at -80 ºC. CB-6644 (MedChemExpress HY-114429), triptolide (AdipoGen AG-CN2-0448-M001), and AZD4573 (Selleckchem S8719) were dissolved in DMSO at 25 mM and stored in aliquots at - 80 ºC. 5-EU (Lumiprobe 2439) was dissolved in sterile-filtered, RNase-free PBS at 100 mM and stored in aliquots at -80 ºC. Palbociclib (AdipoGen CDX-P0590) was dissolved in DMSO at 1 mM and stored in aliquots at -80 ºC. Compounds were diluted from full concentration DMSO stocks into media. Cycloheximide (Sigma Aldrich C7698) was weighed and freshly dissolved on the day of each experiment and diluted to a constant intermediate of 300 µg/mL in 4% DMSO. Leptomycin B (Sigma Aldrich L2913) was supplied in a methanol/water mixture with ratio 7:3, aliquoted and stored at -20 ºC.

For concentration-response experiments in mAC-POLR2A cells using 5-Ph-IAA, serial ten-fold dilutions were made from the highest concentration, with DMSO added to keep final vehicle constant. This series of 5x stocks were added, 20 µL onto 80 µL of media, for final concentrations of 10 nM, 100 nM, 1 µM, or 10 µM 5-Ph-IAA all in 0.1% DMSO, or 0.1% DMSO only. With the exception of this experiment in mAC-POLR2A cells, all other 5-Ph-IAA treatments used 10 µM final concentration.

For palbociclib and cell size experiments, cells were plated in 80 µL of media. 24 hours later, 20 µL palbociclib or vehicle was added at 5x concentration for final concentration of 1 µM in 0.1% DMSO, or 0.1% DMSO only.

For experiments where 5-Ph-IAA treatment was combined with concurrent translation inhibition or nuclear export inhibition, media was exchanged for fresh via a 2x wash, to a final volume of 60 µL. 15 µL of 5-Ph-IAA or vehicle at 6x concentration was added, followed by 15 µL cycloheximide, leptomycin B, or vehicle, for final concentrations of 10 µM 5-Ph-IAA, ± 50 µg/mL cycloheximide, ± 10 ng/mL leptomycin B, in 0.8% DMSO, or 0.8% DMSO only. For experiments where translation inhibition or nuclear export inhibition was performed after pretreatment with 5-Ph-IAA, 11 µL of 5-Ph-IAA or vehicle at 10x concentration was added 24 hours before beginning time-course onto 100 µL for 10 µM 5-Ph-IAA all in 0.1% DMSO, or 0.1% DMSO only. Media was exchanged for fresh via a 3x wash, to a final volume of 60 µL, and 15 µL 5-Ph-IAA or vehicle added at 6x concentrations to return wells to previous treatment. 15 µL cycloheximide, leptomycin B, or vehicle were then added at indicated times, for final concentrations of 50 µg/mL cycloheximide, 10 ng/mL leptomycin B, or vehicle, ± 10 µM 5-Ph-IAA, all in 0.8% DMSO final.

For other fixed cell experiments, cells were plated in 80 µL or 100 µL of media. 5-Ph-IAA or vehicle was added, 20 µL at 5x concentrations or 11 µL at 10x concentrations, at indicated timepoints for final concentrations of 10 µM 5-Ph-IAA in 0.1% DMSO, or 0.1% DMSO only. For live cell experiments (bleach-chase assay and live-cell imaging of fluorescent proteins), 5-Ph-IAA or vehicle was added at 10x concentration in imaging media, 22 µL onto 200 µL, for final concentrations of 10 µM 5-Ph-IAA in 0.1% DMSO, or 0.1% DMSO only.

For FRAP experiments, 5-Ph-IAA or vehicle was added at 10x concentration in imaging media, 40 µL onto 360 µL for final concentrations of 10 nM, 100 nM, 1 µM, or 10 µM 5-Ph-IAA in 0.1% DMSO or 0.1% DMSO only.

In cases where siRNA transfection was followed by 5-Ph-IAA treatment, 14 µL of 5-Ph-IAA or vehicle was added at 6x concentrations on to 70 µL at indicated timepoints for final concentrations of 10 µM 5-Ph-IAA in 0.1% DMSO, or 0.1% DMSO only.

For assaying 5-EU incorporation, cells were initially plated in 100 µL of media. Before beginning treatment time-courses, media was exchanged for fresh via a 2x wash, to a final volume of 60 µL. 15 µL 5-Ph-IAA or vehicle was added at a 5x concentration to a final volume of 75 µL at indicated timepoints for 10 µM 5-PhIAA in 0.1% DMSO or in 0.1% DMSO only. 5-EU was diluted from aliquots of 100 mM into media. 30 minutes before fixation, 15 µL of 6 mM 5-EU in media, with 1x 5-PhIAA or vehicle only to maintain constant concentrations, was added into relevant wells, and plate returned to incubator.

For smFISH experiments, 5-Ph-IAA, triptolide, AZD4573, CB6644, or the corresponding DMSO in media vehicle were added at 10x concentrations at indicated timepoints, 11 µL onto 100 µL, for final concentrations of 10 µM 5-Ph-IAA in 0.1% DMSO or 1 µM triptolide, AZD4573, or CB6644 in 0.004% DMSO. For cell segmentation in smFISH experiments, CF488-conjugated concanavalin A (Biotium 29016) was added in media at 5x concentrations for a final of 50 µg/mL for 10 minutes before fixation.

### Real-time quantitative polymerase chain reaction

3 days after plating, total RNA from all samples was isolated using the Trizol reagent (Invitrogen, 15596026) following the Direct-zol RNA MiniPrep Kit (Zymo, R2052), according to the manufacturer’s protocol. 1 µg extracted RNA for each sample was reverse-transcribed into cDNA using the High Capacity cDNA Reverse Transcription Kit (Applied Biosystems, 4368814), according to manufacturer’s instructions. cDNA was diluted 100 times before qPCR. RT-qPCR reactions were performed in 96-well plates (Bio-Rad, HSP9601) with a Bio-Rad CFX96 Touch Real-Time PCR Detection System. 20 µL qPCR reactions contained 10 µL 2x SsoAdvanced Universal SYBR Green Supermix (Bio-Rad, 1725271), 10 µM primer of each gene-specific primer, and 5 µL diluted cDNA. The following standard thermal profile was used for all qPCR reactions: 95 °C for 30 s, 40 cycles of 95 °C for 15 s, and 60 °C for 15 s. Melting curves were recorded after cycle 40 by heating from 65 °C to 95 °C with an increment of 5 s/step. qPCR data were analyzed in R software using ΔΔCq method. Two standard reference genes, *UBC* and *ACTB*, were used as internal controls for gene expression normalization. For each target gene, ΔCq values were calculated by subtracting the average reference gene Cq from the target gene Cq. To determine relative gene expression, ΔΔCq values were computed by subtracting the mean ΔCq of the scramble control samples from the ΔCq of each experimental sample. Fold changes in gene expression were calculated using the formula 2^(-ΔΔCq)^, with scramble control samples set as the baseline (fold change = 1) (*38*).

### Fixed cell imaging and image processing

Fixed cell imaging was performed using a Nikon Ti2 microscope with Yokogawa CSU-W1 spinning disk using twin Hamamatsu ORCA-Fusion C14440-20UP cameras. For imaging smFISH, a 100x 1.46NA oil immersion objective was used, with 61 z-planes acquired at 250 nm steps. For imaging 7SK FISH a 40x 0.95NA Plan Apo λ objective was used, with 20 z-planes acquired at 1 µm steps. For imaging mCherry-POLR2A abundance in palbociclib or vehicle treated cells a 40x 0.95NA Plan Apo λ objective was used, with 35 z-planes acquired at 600 nm steps. For all other fixed cell experiments (immunofluorescence, fixed fluorescent protein imaging, and 5-EU incorporation), a 20x 0.75NA objective was used, with 20 z-planes acquired at 1 µM steps. DAPI DNA stain was acquired using a 405 nm laser and 450/82 nm filter. Alexa Fluor Plus 488-conjugated secondary antibodies, CF488-conjugated concanavalin A, or mClover3 were acquired using a 488 nm laser and 525/50 nm filter. Alexa Fluor 568-conjugated secondary antibodies, ATTO565 smFISH probes, or mCherry were acquired using a 561 nm laser and either a 617/73 nm filter, or a 590/33 nm narrow-band filter in cases where a far-red stain was also present. Alexa Fluor 647-conjugated secondary antibodies and ATTO633 smFISH probes were acquired using a 640 nm laser and 685/40 nm filter.

Image processing for all fixed cell experiments was performed using custom pipeline written in python, progressing from raw images through to single cell measurements. These were then analysed, summarised, and plotted using python, making use of JupyterLab, numpy, and pandas.

For all experiments except smFISH, z-stacks were maximum projected and corrected for illumination bias across each field of view. Nuclei were segmented in 2D from the DAPI signal using Cellpose3 with built-in model ‘nuclei’. Mean fluorescence intensity and nuclear morphology measurements were calculated using the regionprops function from the scikit-image (*39*) python package. Fluorescence background subtraction was performed using negative control wells within each plate (e.g. secondary-only wells in immunofluorescence, or no 5-EU wells in click chemistry). To combine data from distinct experiments performed on different days, measurements were normalized by dividing all background-subtracted data within each plate by the mean of control wells (e.g. vehicle-treated cells, or scrambled siRNA). Time-matched vehicle control wells, i.e. with vehicle addition at indicated 5-Ph-IAA timepoints, were examined for systematic effects, then averaged and used as t = 0 points for examining 5-Ph-IAA effects.

For smFISH experiments, analysis was performed in 3D on complete z-stacks. For cell segmentation, DAPI and concanavalin A signal was downscaled by ten-fold and segmented with Cellpose3 ‘cyto3’ model. These labels were then returned to original size, eroded, and used as seeds along with the cell-free area for a watershed run on each slice individually, on Sato-filtered concanavalin A signal using scikit-image. Nuclei were similarly segmented by downscaled by ten-fold, segmenting in Cellpose3 using ‘nuclei’ model, returning labels to original size, eroding, and performing a watershed on Roberts-filtered DAPI signal using scikit-image. In both cases, clean-up was performed by removing small objects. Segmented nuclei were then assigned to cells by binarizing and relabelling, such that all volume labelled as nucleus within a cell corresponds to that cell.

Spot detection for smFISH employed the bigfish package (*40*), with ATTO565 or ATTO633 signal Laplacian of Gaussian filtered, after kernel size was set using the object radius calculation function, local maxima detected, then spots thresholded using bigfish functions. Exact detection thresholds for smFISH spots counted as true were set globally for each experiment, using an odd/even control well in which ATTO565 and ATTO633 probes targeted the same transcript.Threshold was set to maximize coincidence between these two channels, with detection efficiency calculated within each experiment and employed as a quality control metric.

### Fluorescence recovery after photobleaching (FRAP) and data analysis

mCherry-POLR2A and mAC/mCherry-POLR2A cells were plated at 8000 cells per well, and mAC-POLR2A cells were plated at 16000 cells per well due to slow growth rates, in cases where siRNAs were not employed in 400 µL of imaging media. The morning of the imaging session, media was exchanged for 360 µL of fresh imaging media additionally containing 1% penicillin/streptomycin. 5-Ph-IAA, 40 µL at 100 µM, for 10 µM final in 0.1% DMSO, or vehicle only was added at staggered time-points such that cells could be collected from each well in the indicated ranges of 5-PhIAA incubation (60 - 90 minutes or 240 – 300 minutes). Wells in 24 hour 5-Ph-IAA treated conditions had addition performed the day before and were immediately returned to constant final 5-Ph-IAA concentrations on media exchange.

All FRAP traces were collected on a Zeiss LSM900 point-scanning confocal with a 63x 1.40NA Plan-Apochromat oil immersion objective at 37ºC and 5% CO_2_. For each cell, an initial image of the whole nucleus was collected. In experiments comparing mCherryPOLR2A, mAC-POLR2A, and mAC-/mCherry-POLR2A cells, images of both mCherry and mClover3 nuclear signal were collected from all cells before FRAP experiment (Fig. S5A). A field of view of 173 square pixels was defined and imaged with a frame interval of 300 ms, and two circular regions were defined with the target nucleus with a diameter of 1.8 µm (18 pixels) and an area of 2.45 µm (255 pixels). In acquired mClover3 FRAP traces, illumination was limited to these defined circular regions to minimise acquisition photobleaching. After a 30 frame baseline, one circular region was bleached with 100% laser power for a fixed number of iterations over approximately 2 s, before imaging continued for a further 60 frames for a total imaging time of approximately 30 s. Control traces were collected under identical conditions from mCherry-POLR2A, mAC-POLR2A, and mAC-/mCherry-POLR2A cells fixed in 4% PFA for 15 minutes (Fig. S5C, S5E).

Raw mean intensity was taken from each circular region of interest, bleached or unbleached, throughout the recording time (Fig. S5B, S5D) and analysed further using python, employing NumPy, pandas, and matplotlib with seaborn. Within each region, fluorescence intensity was individually rescaled to the mean intensity of the 10 s baseline collected. FRAP traces were then normalised with 0% defined by the post-bleach intensity from fixed cells and 100% defined by the unbleached control circular region (Fig. S5C, S5E). FRAP traces collected from fixed cells did not show substantial recovery, reflecting almost completely immobilised fluorescent POLR2A.

Normalised, rescaled FRAP traces were fit to an exponential curve post-bleach using python and scipy, with the recovery plateau value taken as the fraction of mobile POLR2A, and the remaining fraction taken as immobile POLR2A. Fluorescence intensity-adjusted POLR2A amounts in each cell were calculated using the baseline mean intensity of each cell relative to unperturbed biallelic mAC-POLR2A or biallelic mCherry-POLR2A cells.

FRAP traces are shown as mean ± S.D. of all cells within each dataset. Summary plots of amount of POLR2A in immobile or mobile fraction are shown as mean ± S.D. of three separate experiments.

### Western blot

Proteins were separated on 4%-12% or 4%-20% Tris-Glycine gels (Thermo Fisher Scientific) and transferred to nitrocellulose membranes (Amersham). Membranes were blocked in 5% (w/v) skimmed milk in PBS-T (PBS, 0.1% (v/v) Tween-20) for 1 h at room temperature and incubated with primary antibody (in 5% (w/v) skimmed milk in PBST) overnight at 4°C. Primary antibodies against: D8L4Y (Rabbit monoclonal RPB1 (total, N-terminal), Cell Signaling, RRID:AB_2687876); GFP (Rabbit polyclonal, Abcam, ab6556, RRID:AB_ 305564); mCherry (Rabbit polyclonal, Abcam, ab167453, RRID:AB_2571870); CDK9 (Rabbit monoclonal, Cell Signaling, 2316T, RRID:AB_2291505); H3 (Rabbit polyclonal, Abcam, ab18521, RRID:AB_732917); Vinculin (Mouse monoclonal, Sigma, V9131, RRID:AB_477629). Membranes were subjected to 3 x 5 min washes with PBST, incubated in 5% (w/v) skimmed milk in PBST containing HRP-conjugated secondary antibody (anti-mouse, Dako, P044701-2, RRID:AB_2617137; anti-rabbit, Dako, P044801-2, RRID:AB_2617138), and visualised using SuperSignal West Pico PLUS (Thermo Fisher Scientific).

### Chromatin fractionation

For each condition, 3.6 x 10^6^ cells were seeded onto a 10 cm dish. The next day, the cells were scraped in PBS, span down and the pellet was snap-frozen in liquid nitrogen. The frozen pellets were defrosted at room temperature and transferred to ice. 265 μL of soluble extraction buffer (20 mM HEPES-KOH pH=7.5, 150 mM potassium acetate, 1.5 mM MgCl2, 10% v/v glycerol, 0.05% NP-40, with addition of fresh protease inhibitors, phosphatase inhibitors and 2 mM NEM) were used to resuspend each cell pellet, and the suspensions were incubated on ice for 20 minutes. To release the cytosol and nucleoplasm, 20 strokes with a loose micropestle were applied to each sample. The chromatin was pelleted by centrifugation at 1000 rcf at 4ºC for 10 minutes and the supernatant kept as soluble fraction. The pellets were washed with soluble extraction buffer, resuspended in 100 μL of chromatin extraction buffer 1 (125 U/mL of benzonase in 20 mM HEPES-KOH pH=7.5, 1.5 mM MgCl2, 10% glycerol, 150 mM NaCl, 0.05% NP-40, with addition of fresh protease inhibitors, phosphatase inhibitors and 2 mM NEM) and incubated on ice for 30 minutes. Each sample was centrifuged at 20,000 rcf at 4ºC for 10 minutes and the supernatants saved as the first fraction in a new tube. The pellets were resuspended in 50 μL of chromatin extraction buffer 2 (20 mM HEPES-KOH pH=7.5, 1.5 mM MgCl2, 10% glycerol, 3 mM EDTA, 500 mM NaCl, 0.05% NP-40, with addition of protease inhibitors, phosphatase inhibitors and 2 mM NEM) and incubated on ice for 10 minutes with occasional gentle vortexing. The samples were centrifuged at 20,000 rcf at 4ºC for 5 minutes and the supernatant transferred to a new tube as the second chromatin fraction, to which 115 μL of chromatin dilution buffer (20 mM HEPES-KOH pH=7.5, 1.5 mM MgCl2, 10% glycerol, 3 mM EDTA, 0.05% NP-40, with addition of protease inhibitors, phosphatase inhibitors and 2 mM NEM) were added. These diluted second fraction samples were further centrifuged at 20,000 rcf at 4ºC for 5 minutes. The resulting supernatants were combined with the corresponding first chromatin fractions - giving rise to 265 μL of each total chromatin fraction. Protein concentration was determined using Qubit, and equal volumes of soluble and chromatin fractions was analysed by Western blot.

### TT_chem_-seq (nascent RNA-seq)

TT_chem_-seq was performed essentially as described (*22*), with minor modifications (*15*). For each condition, 3.6 x 10^6^ cells were seeded onto a 10 cm dish. The following day, cells were treated with 5-Ph-IAA (*14*) or vehicle, and nascent RNA was *in vivo* labelled with a 1 mM 4sU (Glentham Life Sciences, GN6085) pulse for exactly 15 min. Labelling was stopped by TRIzol (Thermo Fisher Scientific, 15596026) and RNA extracted as described previously (*22*).

As a control for sample preparation, *S. cerevisiae* (strain BY4741, MATa, his3D1, leu2D0, met15D0, ura3D0) 4-thiouracil (4TU)-labelled RNA was spiked into each sample. S. *cerevisiae* were grown in YPD medium overnight, diluted to an OD_600_ of 0.1, and grown to mid-log phase (OD_600_ of 0.8) and incubated with 5 mM 4TU (Sigma-Aldrich, 440736) for 6 min. Yeast can metabolise 4TU and produce 4sU, which gets incorporated into the nascent RNA. Total yeast RNA was extracted using the PureLink RNA Mini kit (Thermo Fisher Scientific, 12183020) following the enzymatic protocol.

For purification of 4sU labelled RNA, 100 μg of human 4sU-labelled RNA was spiked-in with 1 μg of 4sU-labelled *S. cerevisiae* RNA. The 101 μg of RNA (in a total volume of 100 μL) were fragmented by addition of 20 μL freshly made 1 M NaOH and incubated on ice for 20 min. Fragmentation was stopped by addition of 80 μL 1 M Tris pH 6.8 and the samples cleaned up twice with Micro Bio-Spin P-30 Gel Columns (Bio-Rad, 7326223) adding 200 μL of RNA solution per column. The biotinylation of 4sU-residues was carried out in a total volume of 250 μL, containing 10 mM Tris-HCl pH 7.4, 1 mM EDTA and 5 μg MTSEA biotin-XX linker (Biotium, BT90066) for 30 min at room temperature in the dark. The RNA was then purified by phenol-chloroform extraction, denatured by 10 min incubation at 65°C and added to 200 μL of μMACS Streptavidin Microbeads suspension (Milentyl, 130-074-101). The RNA was incubated with the beads for 15 min at room temperature and the mix was applied to a pre-equilibrated μColumn in the magnetic field of a μMACS magnetic separator. Beads were washed twice with wash buffer (100 mM Tris-HCl pH 7.4, 10 mM EDTA, 1 M NaCl and 0.1% Tween20). Biotinylated RNA was eluted twice by addition of 100 mM DTT and cleaned up with RNeasy MinElute kit (QIAGEN, 74204) using 1050 μL of 100% ethanol per 200 μL reaction after addition of 700 μL RLT buffer to also bind short RNA fragments to the silica matrix.

Libraries for RNA sequencing were prepared using the KAPA RNA HyperPrep Kit (KR1350) with modifications. 75 ng of RNA per sample were mixed with FPE Buffer, but fragmentation procedure was omitted and RNA was instead denatured at 65°C for 5 min. The rest of the procedure was performed as recommended by the manufacturer, with the exception of SPRI bead purifications: after adapter ligation, 0.95x and 1x SPRI bead-to-sample volume ratios were used (instead of two rounds of SPRI purification with 0.63x volume ratios). This was done to retain smaller (150-300 bp) cDNA fragments in the library which would otherwise be lost in size selection. The libraries were quality controlled by electrophoresis on a Tapestation system (Agilent), quantified by Qubit (Thermofisher), pooled and sequenced with single end 70 bp reads on a NextSeq2000, with 50,000,000 average reads per sample. Biological triplicates were generated for each condition.

### dxChIP-seq (double-crosslinking chromatin immunoprecipitation and sequencing)

dxChIP-seq was performed step-by-step as we have described (*41*). Two 15 cm dishes were seeded per condition, each containing 8.4 x 10^6^ cells. The following day, cells were treated with DMSO or 5-Ph-IAA before crosslinking. For immunoprecipitation, 50 μL protein G dynabeads (10004D, Fisher Scientific) per sample were pre-washed and pre-coated with 20 μg of antibody (Pol II D8L4Y, RRID:AB_2687876). DNA libraries were prepared with NEBNext Ultra II DNA Library prep kit (E7645L, NEB). The libraries were sequenced with paired end 60 bp reads on a NextSeq2000, with 30,000,000 average reads per sample. Biological triplicates were generated for each condition.

### Computational analysis of ChIP-seq and TT-seq data

#### dxChIP-seq alignment and processing

dxChIP-seq reads were processed predominantly using the nextflow chip-seq workflow (*42*). Briefly, trimmed and quality-filtered with Trim Galore (*43*) by default parameters. Trimmed reads were aligned to the hg38 genome using Bowtie2 (*44*) default parameters. Then, PCR duplicates were marked and removed using Picard (*45*), with further filtering for blacklisted regions (*46*). Each sample replicate BAM files were computed using RPKM-normalized using bedtools genomecov (*47*) then converted into bigwigs using UCSC bedgraphsTobigwigs (*48*). The replicate bigwigs were then merged by samples using UCSC bigwigMerge, with merged coverage bed files generated for downstream analysis via in-house R script

#### dxChIP-seq metagene profiles and quantification

Ensembl-annotated genes (GRCh38.102) were stratified by gene length and split into bins. Genes greater than 1 kb were used for metagene and subsequent analysis, with TSSs and TTSs were defined using a ±500 bp window. Coverage was computed using bedtools (*47*) normalizing to the average signal 5kb upstream of each gene (metagene analysis) or the average signal upstream of genes, with Pol II peaks coverage > 0.5 considered as an actual signal (meta TSS/TTS/gene body quantification). All downstream data processing and visualization was performed using in-house R scripts.

#### Pausing index analysis

The TSS region is defined as -30bp upstream and +300bp downstream of the start of the genes (>1kb) with gene body regions defined as +700bp away from the start of the genes (>1kb). Only genes with Pol II peaks TSS coverage > 0.5 in the 0h timepoint sample of the TSS region were considered as actual signal. The pausing index per gene is calculated using the formula log2(TSS normalised coverage/ gene body normalised coverage), where the coverage of TSS and gene body regions were normalised to the average signal upstream of genes respectively.

#### TT_chem_-seq alignment and processing

TT_chem_-seq reads were trimmed and quality-filtered with Trim Galore using default parameters. Trimmed reads were aligned to the merged hg38 and Saccharomyces cerevisiae (sacCer3) genome using STAR aligner (*49*) with basic two-pass mapping. The aligned BAM files were split by strand orientation. Sample replicates were then normalized using the yeast spike-in reads individually and then computed for coverage using deepTools bamCoverage (*50*). The resulting bigwigs were then merged by strand orientation using UCSC tools bigWigMerge (*48*) .

#### TT_chem_-seq metagene profiles

Methods for TT_chem_-seq metagene profile normalisation were similar to those in Cacioppo et al 2024, excluding the duplicated reads removal step. Spike-in normalised bigwigs were used for mapping, and metagenes were plotted without further background normalisation.

#### TT_chem_-seq quantification and differential expression analysis

For each replicate, read counts for genes were quantified using featureCounts -s 2 parameters (*51*). Pairwise comparisons were performed with DESeq2 (*52*), incorporating spike-in normalization for quantitative comparisons across samples using the function controlGenes. Low-count genes were filtered out by retaining only genes with average normalized counts of at least 10 before differential expression analysis and MA plot density analysis. 0h timepoint samples were used as a baseline for all comparisons (90min vs 0h and 8h vs 0h), significantly regulated genes cut-off was Benjamini-Hochberg adjusted p-value < 0.05) and a minimum log2 fold-change of 1. All downstream data processing and visualization was performed in R using in-house scripts.

## Mathematical modeling

### Nuclear Pol II pools

We model the following Pol II species: Nucleoplasmic (*N*), promoter/TSS-bound (*P*), Gene-body/elongating (*P*).

Allowed transitions between these states are:

*N*↔*P* → *E* (initiation/premature termination, release into elongation)

*E* → *N* (termination)

We model initiation and release into elongation using Michaelis-Menten kinetics, with Michaelis constants *K*_m(*NP*)_ and *K*_m(*PE*)_, respectively. Other transitions are modelled using first-order rate constants *k*_*PN*_ and *k*_*EN*_

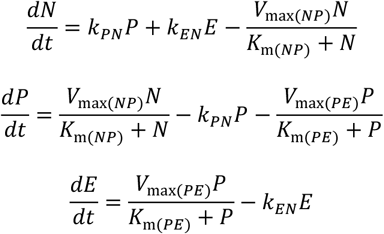

We do not consider polymerase synthesis and decay and require that total polymerase *T* is conserved, *T* = *N* + *P* + *E*, we fit the (steady-state) model to *N, P, E* as a function of *T*, over the range of *T* experimentally accessed. All Pol II concentrations are expressed as fractions of endogenous total Pol II, such that *T* = 1 corresponds to unperturbed conditions. Time is rescaled by fixing the elongation-to-nucleoplasmic return rate *k*_*EN*_ = 1, so that all other rate constants are dimensionless and interpreted relative to this timescale. We also require that 80% of polymerases in state *P* terminate prematurely (rather than proceeding to elongation) when *T* = 1, as previously estimated (*53*). This constraint was implemented by enforcing, at steady state and *T* = 1, that the flux from *P* → *N* constitutes 80% of the total outgoing flux from *P*.

We fit *N* to the unbound concentration of Pol II (free fraction multiplied by mean nuclear intensity) measured by FRAP in mAC-POLR2A upon auxin addition and mCherry-POLR2A in ARMC5 siRNA cells (Fig. 2). We fit *P* using average normalised POLR2A dxChIP-seq coverage over the TSS in both auxin-treated mAC-POLR2A/mCherry-POLR2A monoallelic degron cells (Fig. 3, and in ARMC5 knockout HEK293T cells (*15*); and *E* using pSer2 Pol II immunofluorescence in mAC-POLR2A, mAC-POLR2A/mCherry-POLR2A and mCherry-POLR2A upon ARMC5 depletion (Fig. 1Q). In all cases, total Pol II is derived from immunofluorescence mean nuclear intensity (F12) in the corresponding condition. Control conditions (either vehicle, wild-type, or scrambled siRNA) were normalized to 1. The 5 unconstrained model parameters were fit simultaneously across all data using least-squares regression with cost function equally weighted between the data sources that constrain *N, P*, and *E* (scipy minimize using L-BFGS-B method). The best fit parameter values were *V*_max(*NP*)_ = 2.95, *K*_m(*NP*)_ = 0.298 (0.730*N*_ss_), *k*_*PN*_ = 5.44, *V*_max(*PE*)_ = 0.375, *K*_m(*PE*)_ = 0.0249 (0.0992*P*_ss_), *k*_*EN*_ = 1, where *P*_ss_ and *N*_ss_ are the steady-state values of *P* and *N*, respectively.

### Cytoplasmic Pol II assembly competition

We consider two proteins, *A* (mCherry-POLR2A) and *B* (mAC-POLR2A), that compete for a limiting assembly factor *F* that facilitates their nuclear import.

In the cytoplasm, we consider *A*_*f*_, *B*_*f*_, *F*_*f*_, (free proteins) *AF, BF* (complexes). In the nucleus, we consider *A*_*n*_, *B*_*n*_ (nuclear proteins).

We assume:

1. Both proteins bind the assembly factor in the cytoplasm with equal affinity.
2. Only *AF* and *BF* complexes result in nuclear import of *A* and *B*, each at the same rate.
3. Upon import, *F* is released from *AF* and *BF* complexes and re-enters the cytoplasmic pool.
4. *A* and *B* degrade in both compartments, with different rates.
5. When bound to *F, A* and *B* are protected from degradation.
6. Both proteins are synthesized at the same constant rate.

We do not model synthesis or decay of *F*, but require its conservation,

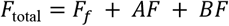

Assembly and nuclear import are captured with the following transitions,

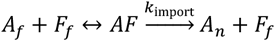

Where *k*_on_ and *k*_off_ are the binding and unbinding rates and *k*_import_ is the import rate. *A*_*f*_, *B*_*f*_ are synthesized with rate *k*_*s*_,

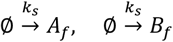

And degraded/diluted with rates,

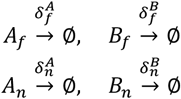

The ordinary differential equation (ODE) governing *AF* is,

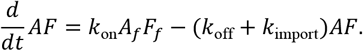

Under the quasi-steady-state assumption, 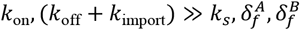, this simplifies to,

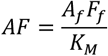

where 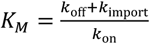 is the Michaelis constant. Conservation of *F* gives,

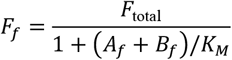

And the remaining ODEs can then be written as,

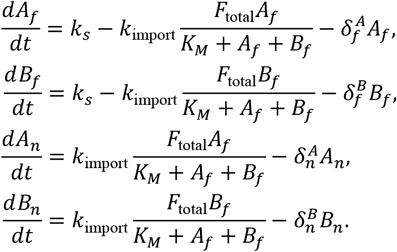

Competition between *A* and *B* is governed by the Michaelis constant *K*_*M*_, with high competition when *K*_*M*_ ≪ *A*_*f*_ + *B*_*f*_.

In mAC-/mCherry-POLR2A monoallelic degron cells, mClover intensity is, on average, 41.53% of that seen in biallelic mAC-POLR2A cells, while mCherry intensity is 59.94% of that seen in biallelic mCherry cells, and immunofluorescence indicates the same total nuclear Pol II levels. We assume that this difference is driven by increased turnover of mAC-POLR2A compared to mCherry POLR2A. For mCherry-POLR2A, we use the measured removal rate for mCherry-POLR2A in bleach chase 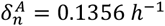 which is equivalent to a protein half-life of 7 h plus dilution due to cell growth (Fig. S10A). Due to the assumed increase in basal degradation of mAC-POLR2A, we set 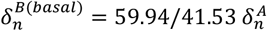, to achieve the experimentally observed ratio. To simulate auxin-induced mAC-POLR2A degradation, we use 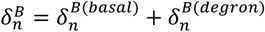 in the nucleus, where 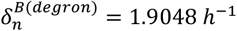 (fit to the loss of nuclear mClover intensity seen upon 5-Ph-IAA addition) and, in the cytoplasm, 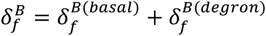. For simplicity, we set 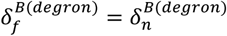, which means that 5-Ph-IAA induced decay rate can occur equally in both compartments. Cytoplasmic decay rates of POLR2A before incorporation into Pol II holoenzyme 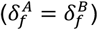 are unknown and were fit as a free parameter.

The following parameter values were obtained by fitting data for plotting in Fig. 5H-I.

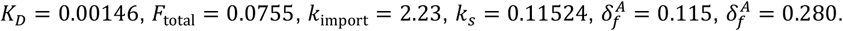

To simulate ARMC5 depletion (Fig. 5I), we set 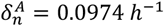 and 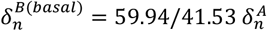, keeping all other parameters constant. For *A* (mCherry-POLR2A), this is equivalent to a protein half-life of 11.4h plus dilution due to cell growth, as previously measured in biallelic mCherry-POLR2A cells using bleach-chase (*15*).

### Nuclear feedback-sensing

As a hypothetical feedback-sensing model for POLR2A synthesis rate adaptation when nuclear POLR2A levels are reduced, we consider the following model,

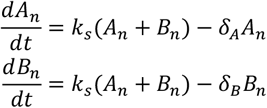

where *A*_*n*_ is nuclear mCherry-POLR2A and *B*_*n*_ is nuclear mAC-POLR2A and *δ*_*A*_, *δ*_*B*_ are nuclear decay rate constants. The synthesis rate *k*_*s*_ (*A*_*n*_ + *B*_*n*_) captures all cytoplasmic processes resulting in appearance of Pol II in the nucleus (e.g. translation, assembly, and nuclear import) and is hypothesized, for the sake of argument, to depend on the total nuclear POLR2A level,

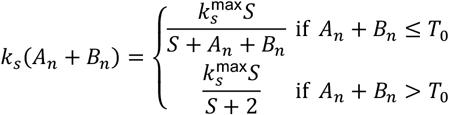

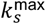 is the maximal synthesis rate, *S* is the feedback sensitivity, *δ*_*A*_, *δ*_*B*_ are degradation rate constants, and *T*_0_ = 0.5994 + 0.4153 = 1.0147 is the steady-state value of total nuclear Pol II. The model is asymmetric (responding only to decreases in nuclear POLR2A but not increases) because we previously found that when nuclear POLR2A levels are elevated through reduced degradation (siRNA-mediated ARMC5 depletion), POLR2A protein synthesis is not reduced to compensate (*15*). All protein decay rates, 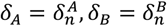 are identical to those defined in the cytoplasmic Pol II assembly competition model, including for ARMC5 knockdown and 5-Ph-IAA-induced degradation. Many combinations of 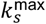 and *S* gave satisfactory fits to 5-Ph-IAA and washout data. In Fig. 5, we used 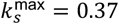 and *S* = 0.3.

