## Supplemental Figures for "Hierarchical buffering of cellular RNA polymerase II pools maintains transcriptional homeostasis"

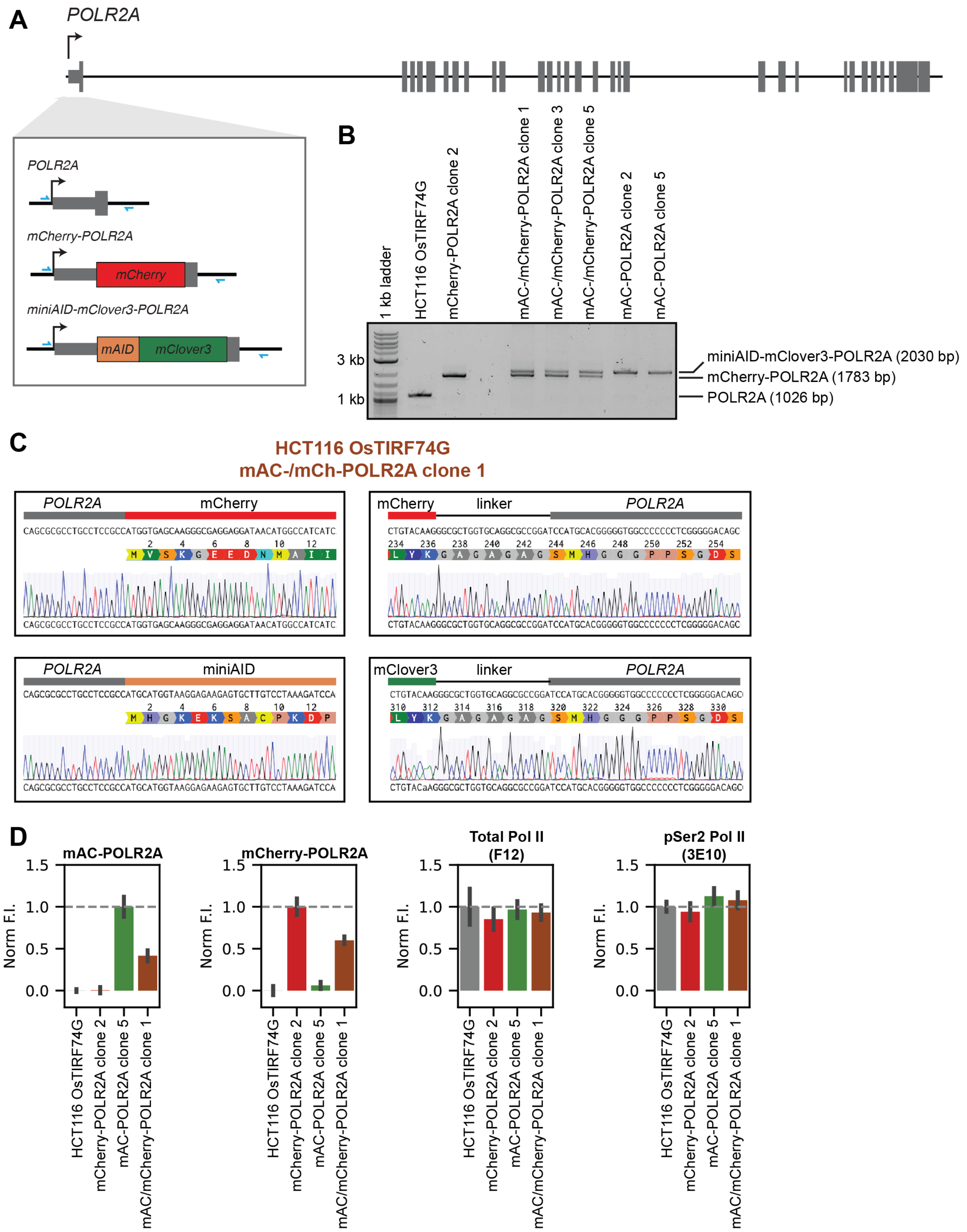


**Supplemental Figure 1. Generation of mAC-/mCherry-POLR2A and biallelic mAC-POLR2A knock-in cell lines**

(A) Genomic structure of the *POLR2A* gene, showing insertion site for mCherry or miniAID-mClover3 fusion protein. Positions of primers used for PCR genotyping shown in blue.

(B) PCR genotyping of parental HCT116 OsTIRF74G cells, biallelic mCherry-POLR2A knock-in cells described previously, mAC-/mCherry-POLR2A monoallelic degron knock-in cells, and biallelic mAC-POLR2A knock-in cells.
(C) Sanger sequencing of edited mCherry-POLR2A and miniAID-mClover3-POLR2A from genomic DNA extracted from mAC-/mCherry-POLR2A clone 1 cells.
(D) Quantification of mClover fluorescence, mCherry fluorescence, total Pol II immunofluorescence and pSer2 Pol II (3E10) immunofluorescence of parental HCT116 OsTIRF74G, mCherry-POLR2A knock-in cells, mAC-/mCherry-POLR2A monoallelic degron knock-in cells, and biallelic mAC-POLR2A knock-in cells, unperturbed or treated with vehicle only. Shown as mean ± S.D. of replicate wells collected in three separate experiments. Note that parental cells of mCherry-POLR2A clone 2 are HCT116, not HCT116 OsTIRF74G.

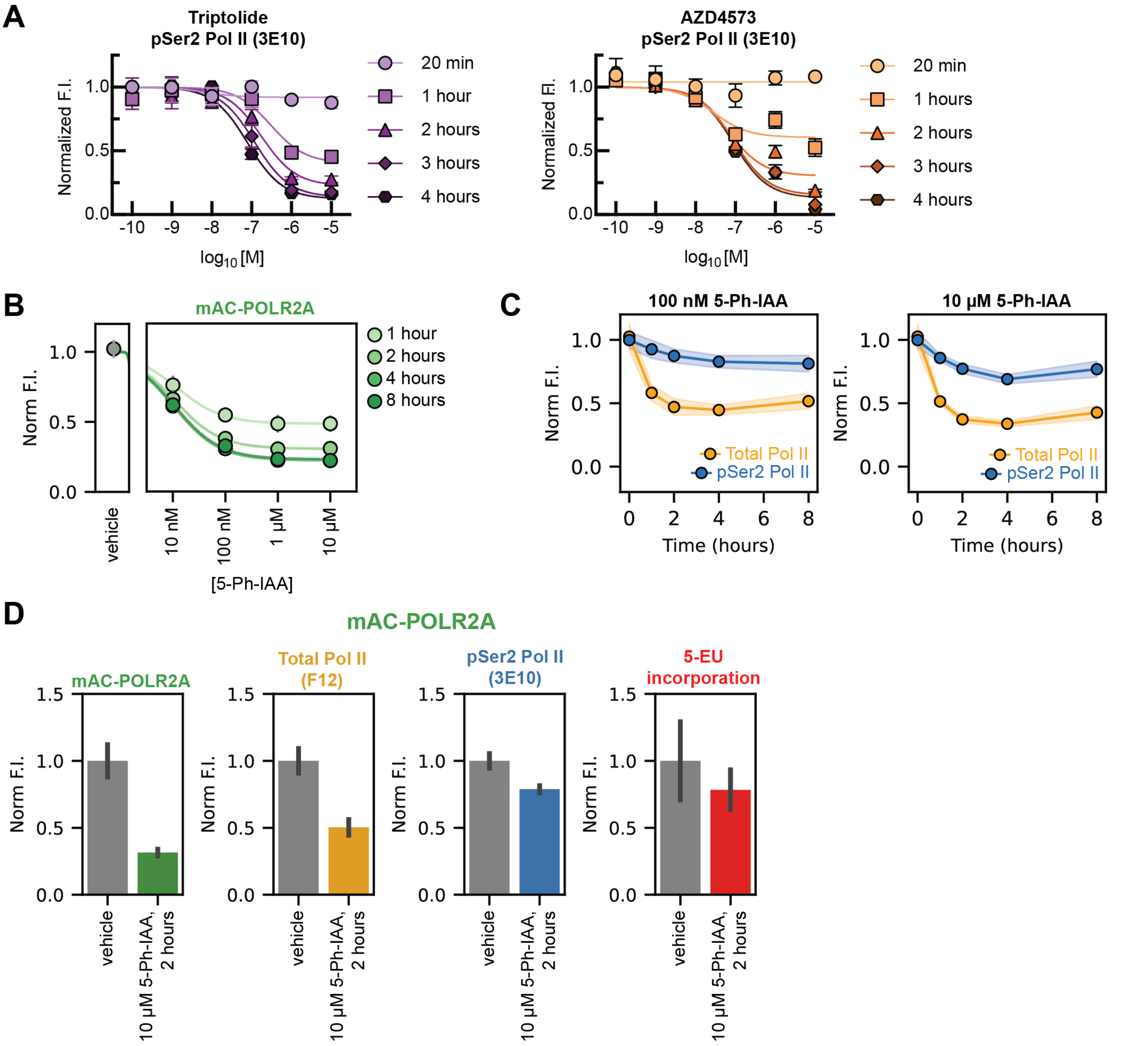


**Supplemental Figure 2. Extended data for mAC-POLR2A biallelic knock-in treated with various 5-Ph-IAA concentrations**

(A) Concentration-response curve for normalized pSer2 Pol II in wild-type HCT116 cells following treatment with either triptolide or AZD4573 for various times. Data shown as mean ± range of two experiments. Mean data fit to a Hill equation with a fixed slope of 1, and top value fixed to vehicle-treated mean of 1.

(B) Concentration-response curve for normalized mClover signal in mAC-POLR2A cells treated with 10 nM, 100 nM, 1 µM, or 10 µM for 1, 2, 4, or 8 hours. Mean data fit to a Hill equation with a fixed slope of 1, and top value fixed to vehicle-treated mean of 1. Points shown as mean ± S.D. of replicate wells from two separate experiments. Note error bars are narrower than points in some cases and not visible.

(C) Total Pol II (F12) and pSer2 Pol II (3E10) immunofluorescence in mAC-POLR2A cells treated with either 100 nM or 10 µM of 5-Ph-IAA. Points shown as mean ± S.D. of replicate wells from two separate experiments.

(D) mClover intensity, total Pol II (F12) immunofluorescence, pSer2 Pol II (3E10) immunofluorescence, and 5-EU incorporation in mAC-POLR2A cells treated with vehicle or 10 µM 5-Ph-IAA. Bars shown as mean ± S.D. of replicate wells from three separate experiments.

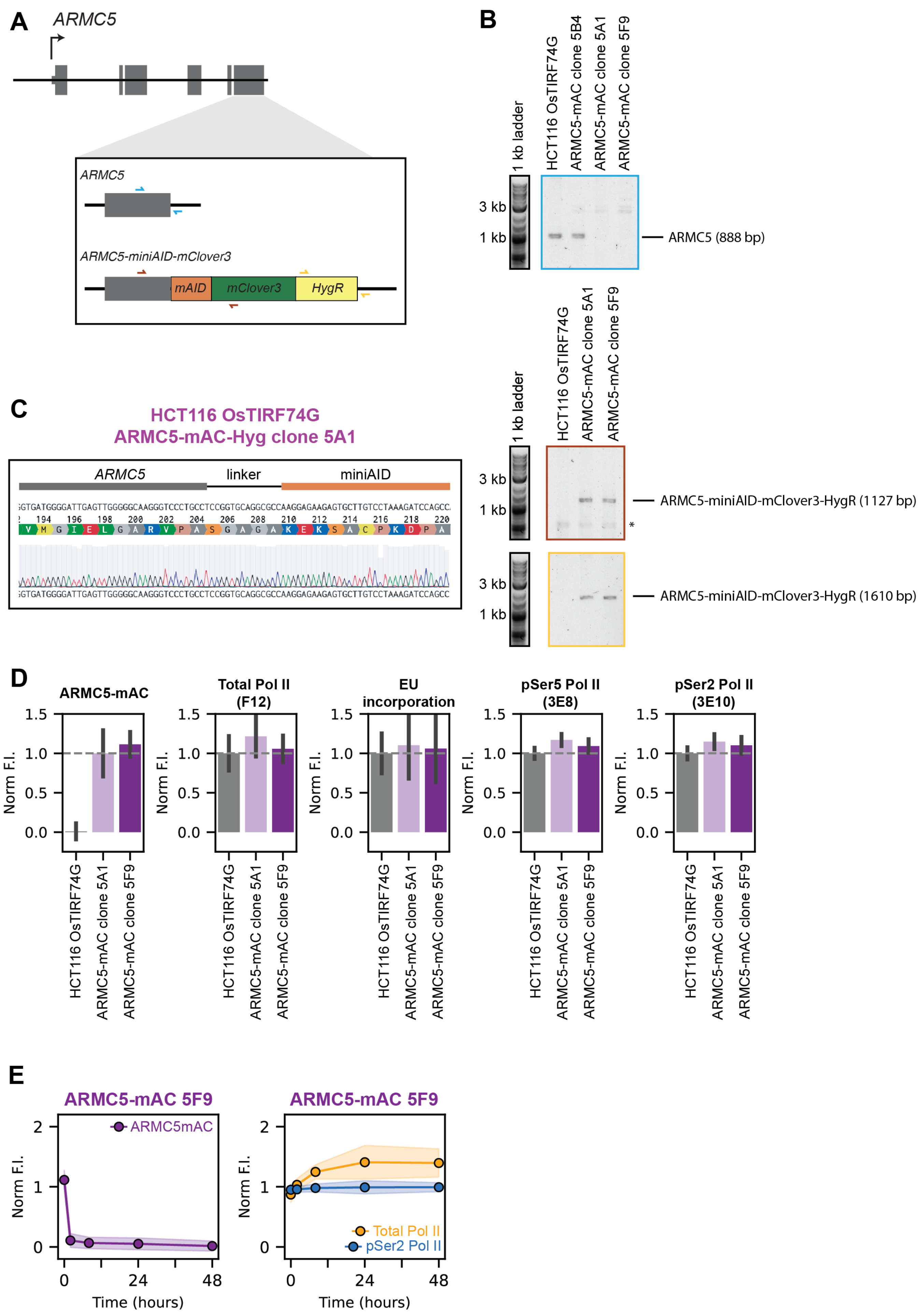


**Supplemental Figure 3. Generation of ARMC5-mAC knock-in cell lines**

(A) Genomic structure of the *ARMC5* gene, showing insertion site for miniAID-mClover3 fusion protein and HygR resistance marker. Positions of primers used for PCR genotyping shown in blue, red, and yellow.
(B) PCR genotyping of parental HCT116 OsTIRF74G cells and ARMC5-mAC knock-in cells.
(C) Sanger sequencing of edited miniAID-mClover3-POLR2A from genomic DNA extracted from ARMC5-mAC clone 5A1 cells.
(D) mClover3 fluorescence, 5-EU incorporation, and total Pol II, pSer5 Pol II (3E8) and pSer2 Pol II (3E10) immunofluorescence of parental HCT116 OsTIRF74G and biallelic ARMC5-mAC knock-in cells, unperturbed or treated with vehicle only. Shown as mean ± S.D. of replicate wells collected in two separate experiments.
(E) ARMC5-mAC mClover3 fluorescence, total Pol II immunofluorescence, and pSer2 Pol II (3E10) immunofluorescence of ARMC5-mAC clone 5F9 cells. Shown as mean ± S.D. of replicate wells collected in two separate experiments.

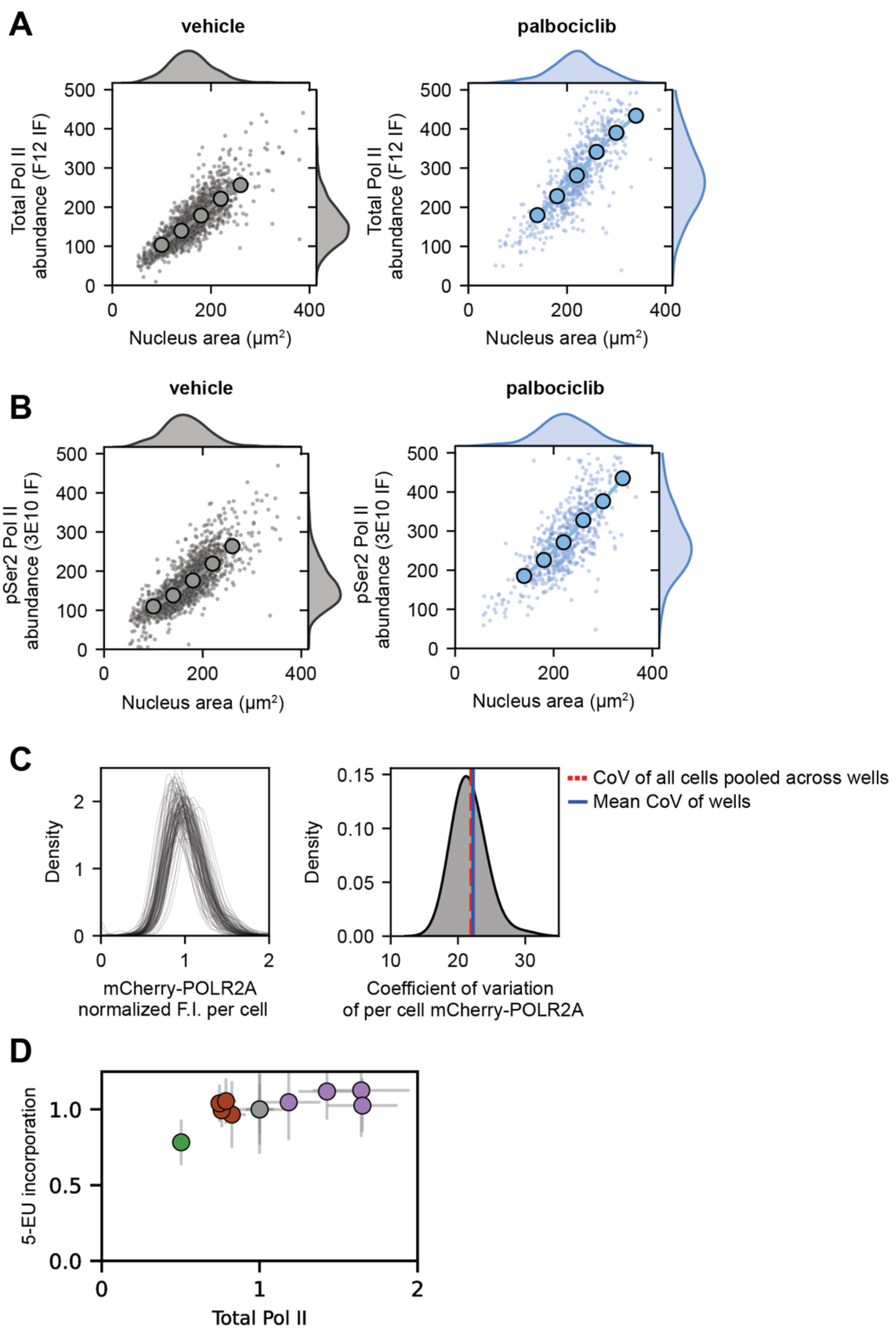


**Supplemental Figure 4. Single-cell Pol II measurements.**

(A) Abundance (integrated nuclear intensity) of total Pol II (F12) as estimated via immunofluorescence in mCherry-POLR2A cells treated with either vehicle or palbociclib compared to nucleus area. Points show one-tenth subsample of entire dataset. Data aggregated across replicates in two separate experiments.

(B) Abundance of pSer2 Pol II (3E10) as estimated via immunofluorescence in mCherry-POLR2A cells treated with either vehicle or palbociclib compared to nucleus area. Points show one-tenth subsample of entire dataset. Data aggregated across replicates in two separate experiments.

(C) Distribution of normalized mCherry-POLR2A fluorescence intensity within individual replicate wells across two experiments shown as single lines, and distribution of the coefficient of variance of normalized mCherry-POLR2A calculated within each replicate well.

(D) Data from Figure 1O and Supplemental Figure 2D showing relationship between Pol II abundance (via F12 immunofluorescence) and 5-EU incorporation into nascent RNA.

Data is shown as mean ± S.D. of replicate wells collected in 2-3 separate experiments.

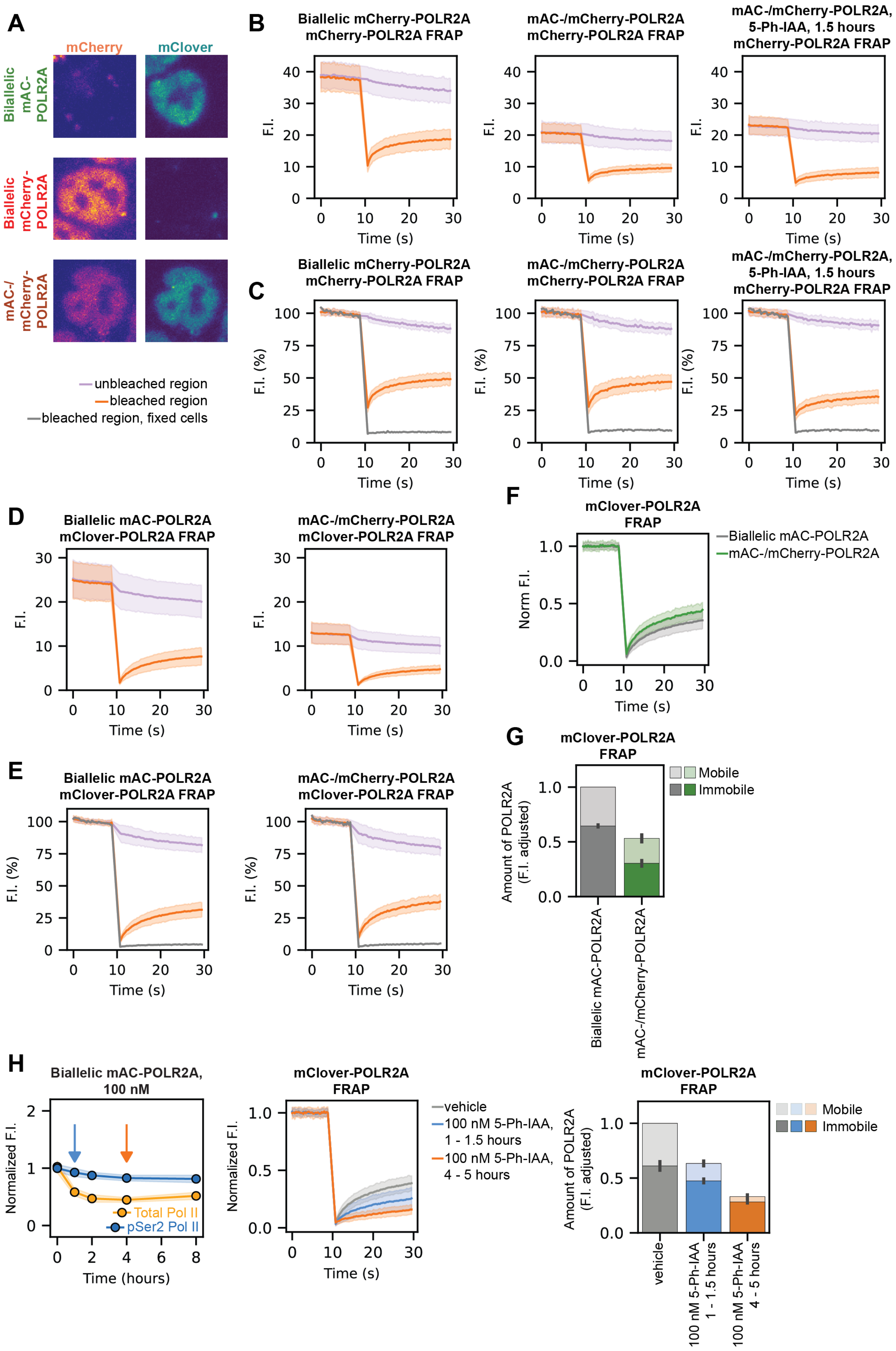
A
mCherry
mClover
Biallelic mAC-POLR2A
Biallelic mCherry-POLR2A
mAC-/ mCherry-POLR2A
unbleached region
bleached region
bleached region, fixed cells
B
Biallelic mCherry-POLR2A mCherry-POLR2A FRAP
mAC-/mCherry-POLR2A mCherry-POLR2A FRAP
mAC-/mCherry-POLR2A, 5-Ph-IAA, 1.5 hours mCherry-POLR2A FRAP
F.I.
Time (s)
C
F.I. (%)
D
Biallelic mAC-POLR2A mClover-POLR2A FRAP
mAC-/mCherry-POLR2A mClover-POLR2A FRAP
E
F
mClover-POLR2A FRAP
Norm F.I.
Biallelic mAC-POLR2A
mAC-/mCherry-POLR2A
G
Amount of POLR2A (F.I. adjusted)
Mobile
Immobile
H
Biallelic mAC-POLR2A, 100 nM
Normalized F.I.
Time (hours)
Total Pol II
pSer2 Pol II
vehicle
100 nM 5-Ph-IAA, 1 - 1.5 hours
100 nM 5-Ph-IAA, 4 - 5 hours

**Supplemental Figure 5. Fluorescence recovery after photobleaching.**
(A) Representative images of those taken of mCherry-POLR2A, mAC-POLR2A, and mAC-/mCherry-POLR2A nuclei before FRAP experiments.
(B) mCherry FRAP traces of mCherry-POLR2A and mAC-/mCherry-POLR2A, treated with either vehicle or 5-Ph-IAA for 1.5 hours, as mean raw fluorescence intensity.
(C) mCherry FRAP traces as in B but baseline corrected, with fixed cell data shown in grey.
(D) mClover FRAP traces of mAC-POLR2A and mAC-/mCherry-POLR2A as mean raw fluorescence intensity.
(E) mClover FRAP traces as in D but baseline corrected, with fixed cell data shown in grey.
(F) Normalized kinetic mClover3 FRAP data of mAC-POLR2A and mAC-/mCherry-POLR2A cells.
(G) Amount of mClover3-POLR2A mobile or immobile estimated by FRAP in mAC-POLR2A and mAC-/mCherry-POLR2A cells, adjusted for fluorescence intensity.
(H) mClover-POLR2A FRAP in mAC-POLR2A cells treated with 100 nM 5-Ph-IAA for 1-1.5 hours, or 4-5 hours, and summary showing amount of POLR2A mobile or immobile.
All kinetic FRAP traces shown as mean ± S.D. of 45 cells collected in three separate experiments, all summary plots of mobile & immobile POLR2A shown as mean ± S.D. of separate experiments.

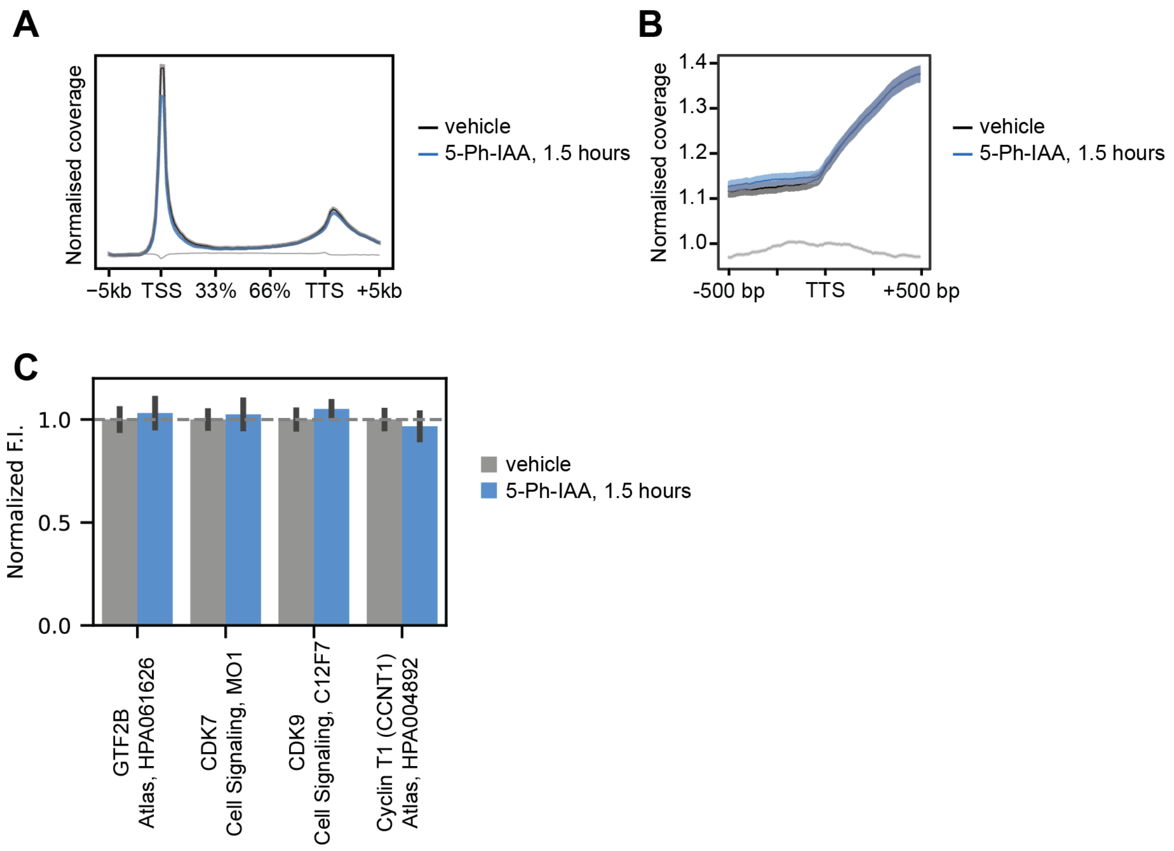


**Supplemental Figure 6.**
(A) Total Pol II dxChIP-seq metagene profile in mAC-/mCherry-POLR2A cells treated with either vehicle or 10 μM 5-Ph-IAA for 1.5 hours.
(B) Same as A but specifically TTS ± 500 bp range.
(C) Immunofluorescence for key Pol II transcription regulators in mAC-/mCherry-POLR2A cells treated with either vehicle or 10 μM 5-Ph-IAA for 1.5 hours.

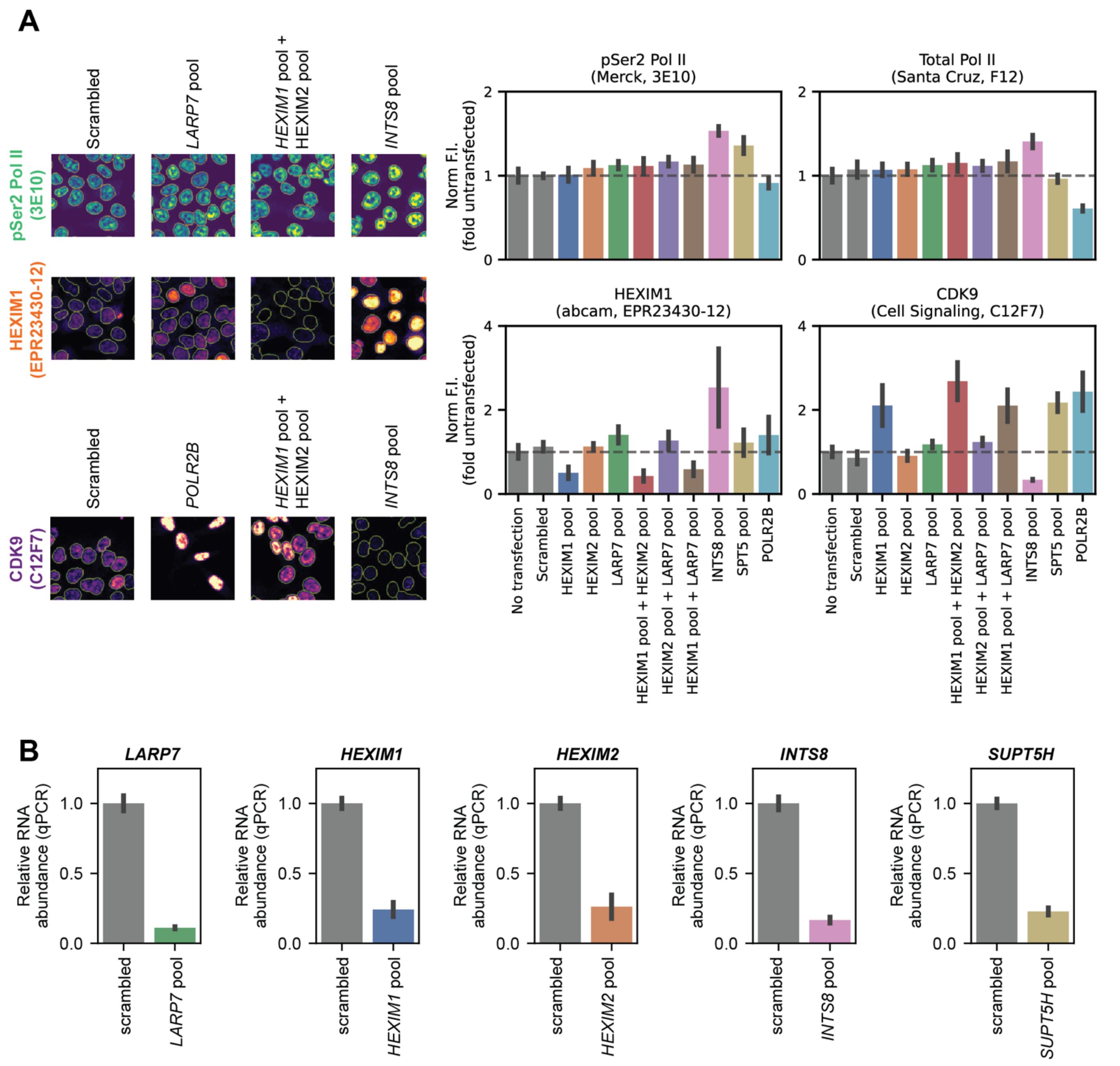


**Supplemental Figure 7.**
(A) Representative images of pSer2 Pol II (3E10), HEXIM1, and CDK9 immunofluorescence in HCT116 cells transfected with either scrambled siRNA, or those targeting LARP7, HEXIM1 plus HEXIM2, INTS8, or POLR2B. Quantification of total Pol II (F12), pSer2 Pol II (3E10), HEXIM1, and CDK9 immunofluorescence in HCT116 cells with various siRNA knockdown. Error bars shown mean ± S.D. of replicate wells collected in two separate experiments.
(B) RT-qPCR validation of *LARP7*, *HEXIM1*, *HEXIM2*, *INTS8*, and *SUPT5H* (SPT5) knock-down by targeted pools.

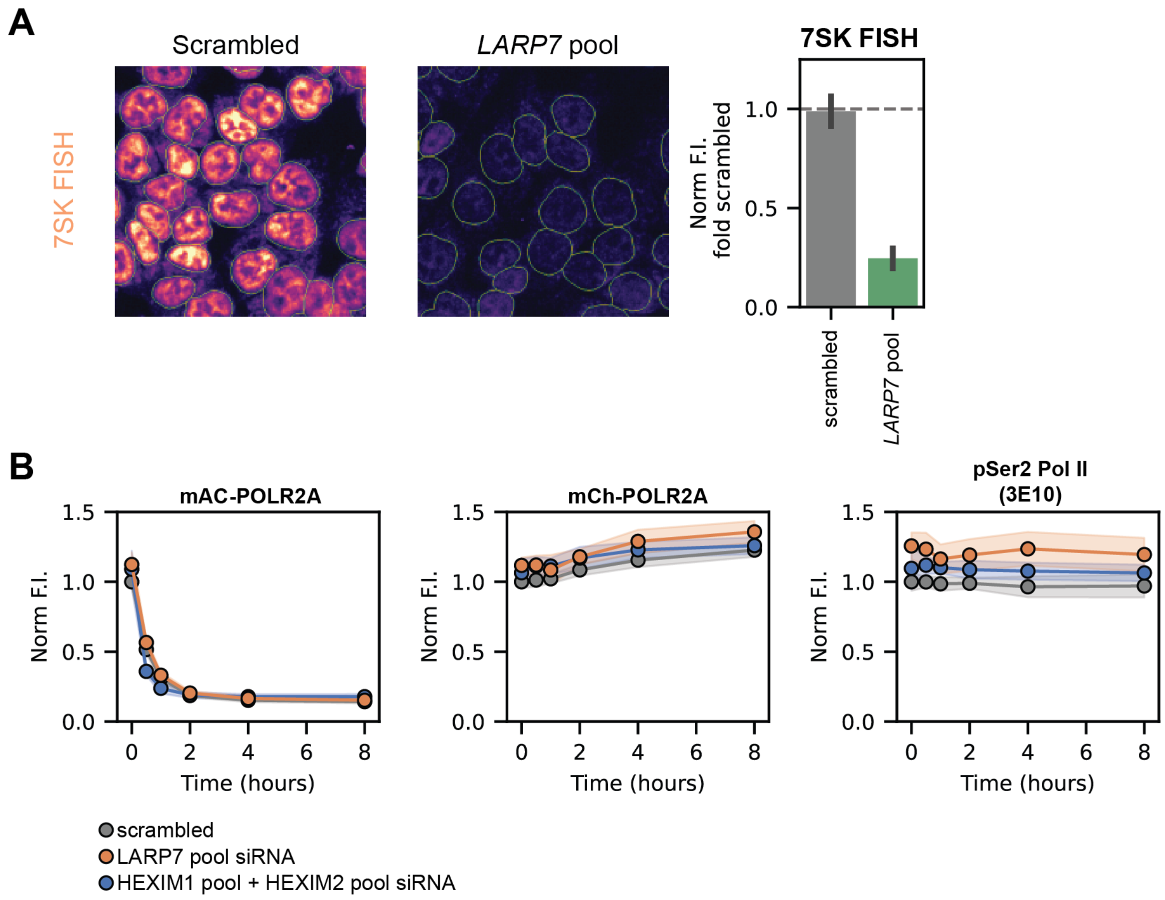


**Supplemental Figure 8.**

(A) Fluorescence in-situ hybridization (FISH) for 7SK ncRNA using ATTO647N-labelled probes, following transfection with either scrambled or LARP7 siRNAs. Quantification shows mean ± S.D. of replicate wells across three separate experiments. Normalization performed with 1 as mean of scrambled siRNA wells, and 0 as wells with no probes added.

(B) mClover3-POLR2A, mCherry-POLR2A, or pSer2 Pol II (3E10) immunofluorescence of mAC-/mCherry-POLR2A cells following transfection with scrambled, *LARP7*-targeted, or *HEXIM1*- with *HEXIM2*-targeted siRNA pools, and 5-Ph-IAA treatment at various timepoints. Quantification shows mean ± S.D. of replicate wells collected in two separate experiments. Significant effect of siRNA knockdown condition on pSer2 levels ($F = 139.9$, $p < 0.001$), but no significant effect of 5-Ph-IAA treatment on pSer2 ($F = 2.2$, $p = 0.14$) nor a significant interaction between siRNA and 5-Ph-IAA treatment ($F = 0.9$, $p = 0.39$) as tested by two-way ANOVA at two hours of 5-Ph-IAA or vehicle treatment (minimum of total Pol II levels).

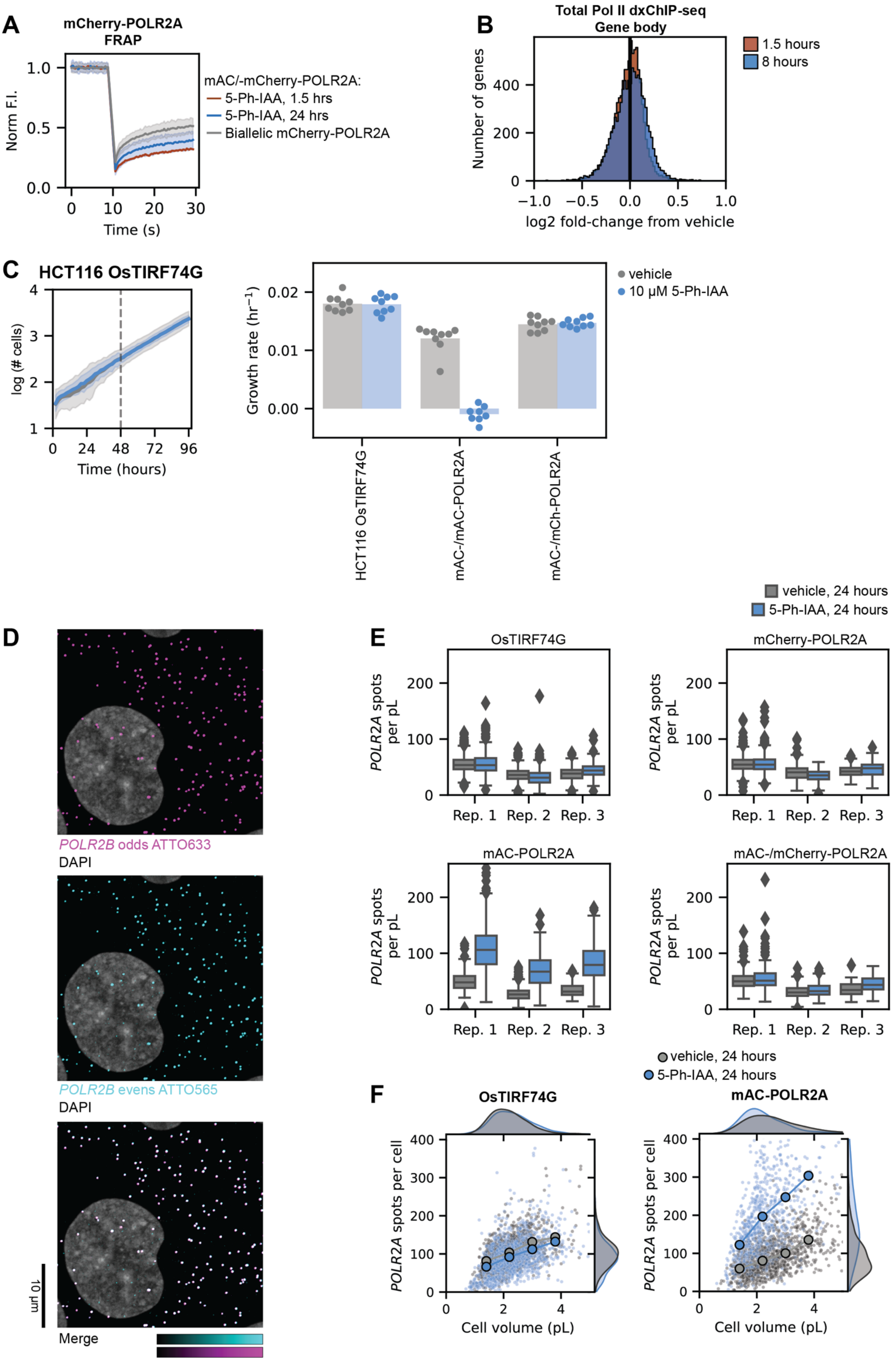

A
mCherry-POLR2A FRAP
Norm F.I.
Time (s)
mAC/-mCherry-POLR2A:
5-Ph-IAA, 1.5 hrs
5-Ph-IAA, 24 hrs
Biallelic mCherry-POLR2A
B
Total Pol II dxChIP-seq
Gene body
1.5 hours
8 hours
Number of genes
log2 fold-change from vehicle
C
HCT116 OsTIRF74G
log (# cells)
Time (hours)
Growth rate (hr$^{-1}$)
vehicle
10 µM 5-Ph-IAA
HCT116 OsTIRF74G
mAC-/mAC-POLR2A
mAC-/mCh-POLR2A
D
POLR2B odds ATTO633
DAPI
POLR2B evens ATTO565
DAPI
Merge
10 µm
E
vehicle, 24 hours
5-Ph-IAA, 24 hours
OsTIRF74G
mCherry-POLR2A
mAC-POLR2A
mAC-/mCherry-POLR2A
POLR2A spots per pL
Rep. 1
Rep. 2
Rep. 3
F
vehicle, 24 hours
5-Ph-IAA, 24 hours
OsTIRF74G
mAC-POLR2A
POLR2A spots per cell
Cell volume (pL)

**Supplemental Figure 9.**
(A) mCherry-FRAP in mAC-/mChPOLR2A cells treated with 5-Ph-IAA for 24 hours, or biallelic mCherry-POLR2A cells. Lines shown mean ±S.D. of 45 cells collected in three separate experiments. Data of 1.5 hours 5-Ph-IAA from Figure 2A included for comparison.
(B) log2 fold-change of gene body coverage in total Pol II dxChIP-seq following treatment with 5-Ph-IAA for 1.5 hours or 8 hours, compared to vehicle.
(C) Total number of cells quantified over 96 hours of HCT116 OsTIRF74G growth, treated with either vehicle or 5-Ph-IAA. Line shows mean ± S.D. of replicate wells collected in two experiments. Growth rates of parental HCT116 OsTIRF74G, mAC-POLR2A, and mAC-/mCherry-POLR2A cells treated with either vehicle or 5-Ph-IAA over 96 hours. Linear fit performed to log-transformed number of cells, from 48 – 96 hours. Points show individual wells collected in two separate experiments.
(D) Example images of *POLR2B* odd and even probe sets used to determine hybridization efficiency and set thresholds using odd/even wells within each individual smFISH experiment.
(E) Effect of vehicle or 5-Ph-IAA treatment for 24 hours on *POLR2A* transcript concentration in HCT116 OsTIRF74G, mCherry-POLR2A, mAC-POLR2A, or mAC-/mCherry-POLR2A cells.
(F) Comparison of volume-dependence of *POLR2A* transcript abundance following 5-Ph-IAA treatment in HCT116 OsTIRF74G, or mAC-POLR2A cells.

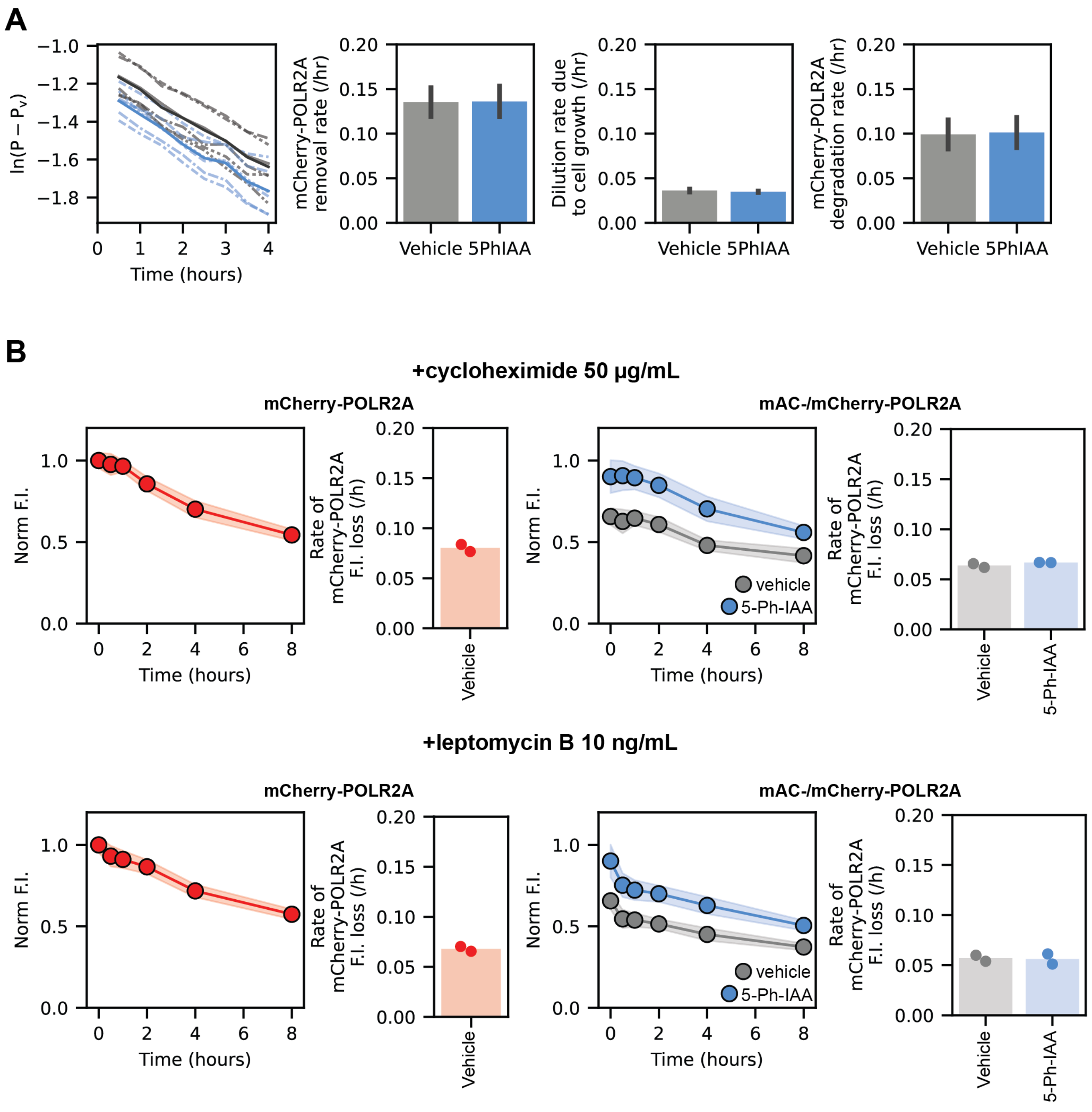


**Supplemental Figure 10.**
(A) Supplemental data for bleach-chase assay showing calculation of Pol II degradation rate, from removal rate and dilution due to cell growth, in mAC-/mCherry-POLR2A cells. Bars shown mean ± S.D. of six replicate wells collected in three experiments.
(B) Loss of nuclear mCherry-POLR2A in biallelic mCherry-POLR2A cells, or mAC-/mCherry-POLR2A cells treated with either vehicle or 5-Ph-IAA for 24 hours, following cycloheximide or leptomycin B treatment. Lines show mean ± S.D. of replicate wells collected in two separate experiments. Points show results of fitting to mean mCherry-POLR2A fluorescence decay of each individual experiment.

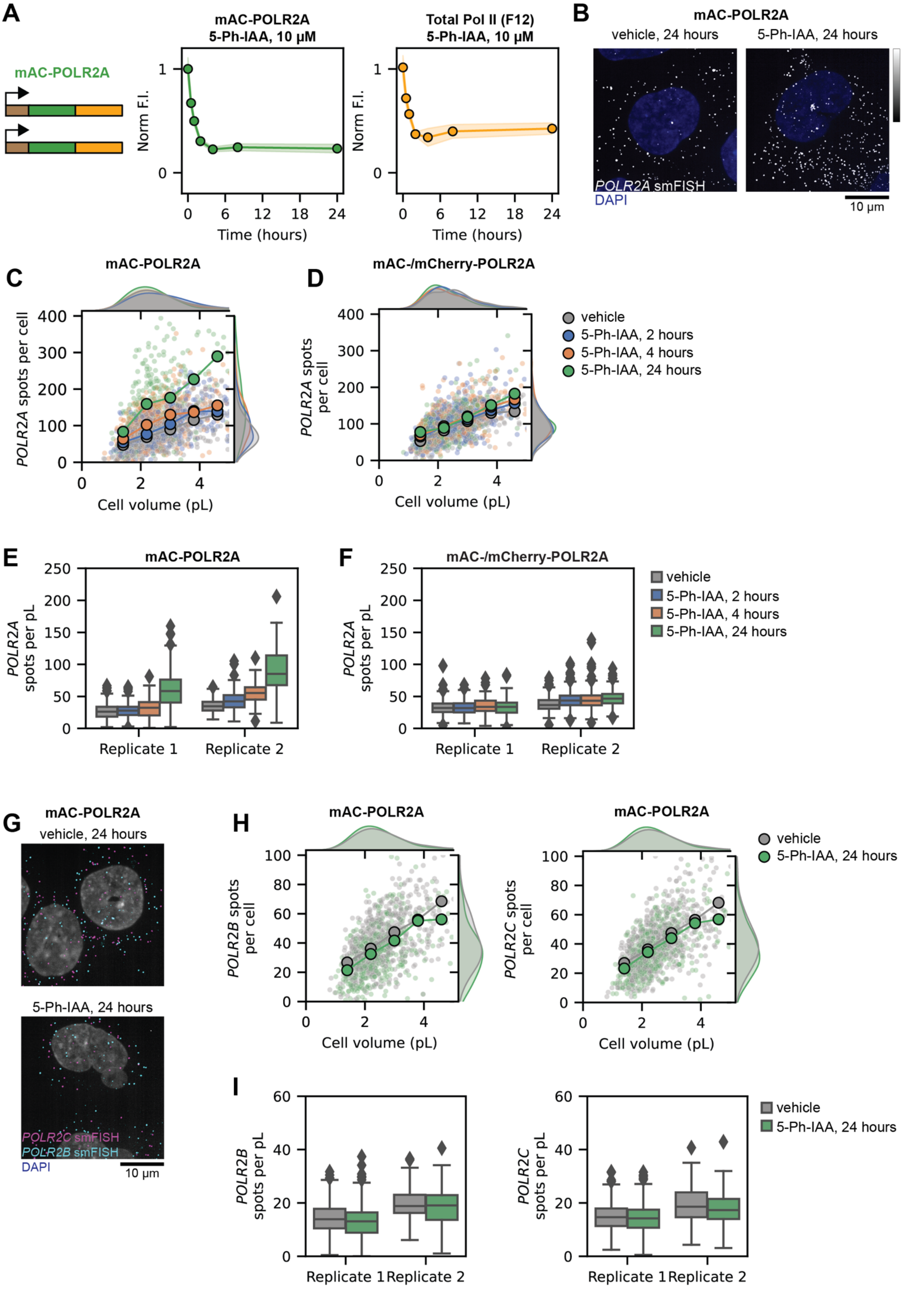


**Supplemental Figure 11.**

(A) Loss of mAC-POLR2A, and total Pol II (F12) immunofluorescence, in mAC-POLR2A cells treated with 5-Ph-IAA at multiple timepoints. Points show mean ± S.D. of replicate wells collected in three separate experiments.
(B) Example images of smFISH for *POLR2A* in mAC-POLR2A cells treated with 5-Ph-IAA or vehicle for 24 hours.
(C) smFISH of *POLR2A* transcript volume-dependence in mAC-POLR2A cells treated with 5-Ph-IAA for 2, 4, or 24 hours.
(D) smFISH of *POLR2A* transcript volume-dependence in mAC-/mCherry-POLR2A cells treated with 5-Ph-IAA for 2, 4, or 24 hours.
(E) Summary of *POLR2A* transcripts per volume across replicate experiments in mAC-POLR2A cells treated with 5-Ph-IAA for 2, 4, or 24 hours.
(F) Summary of *POLR2A* transcripts per volume across replicate experiments in mAC-/mCherry-POLR2A treated with 5-Ph-IAA for 2, 4, or 24 hours.
(G) Example image of smFISH for *POLR2B* and *POLR2C* in mAC-POLR2A cells treated with 5-Ph-IAA or vehicle for 24 hours.
(H) smFISH of *POLR2B* and *POLR2C* transcript volume-dependence in mAC-POLR2A cells treated with 5-Ph-IAA for 2, 4, or 24 hours.
(I) Summary of *POLR2B* and *POLR2C* transcripts per volume across replicate experiments in mAC-POLR2A cells treated with 5-Ph-IAA for 2, 4, or 24 hours.

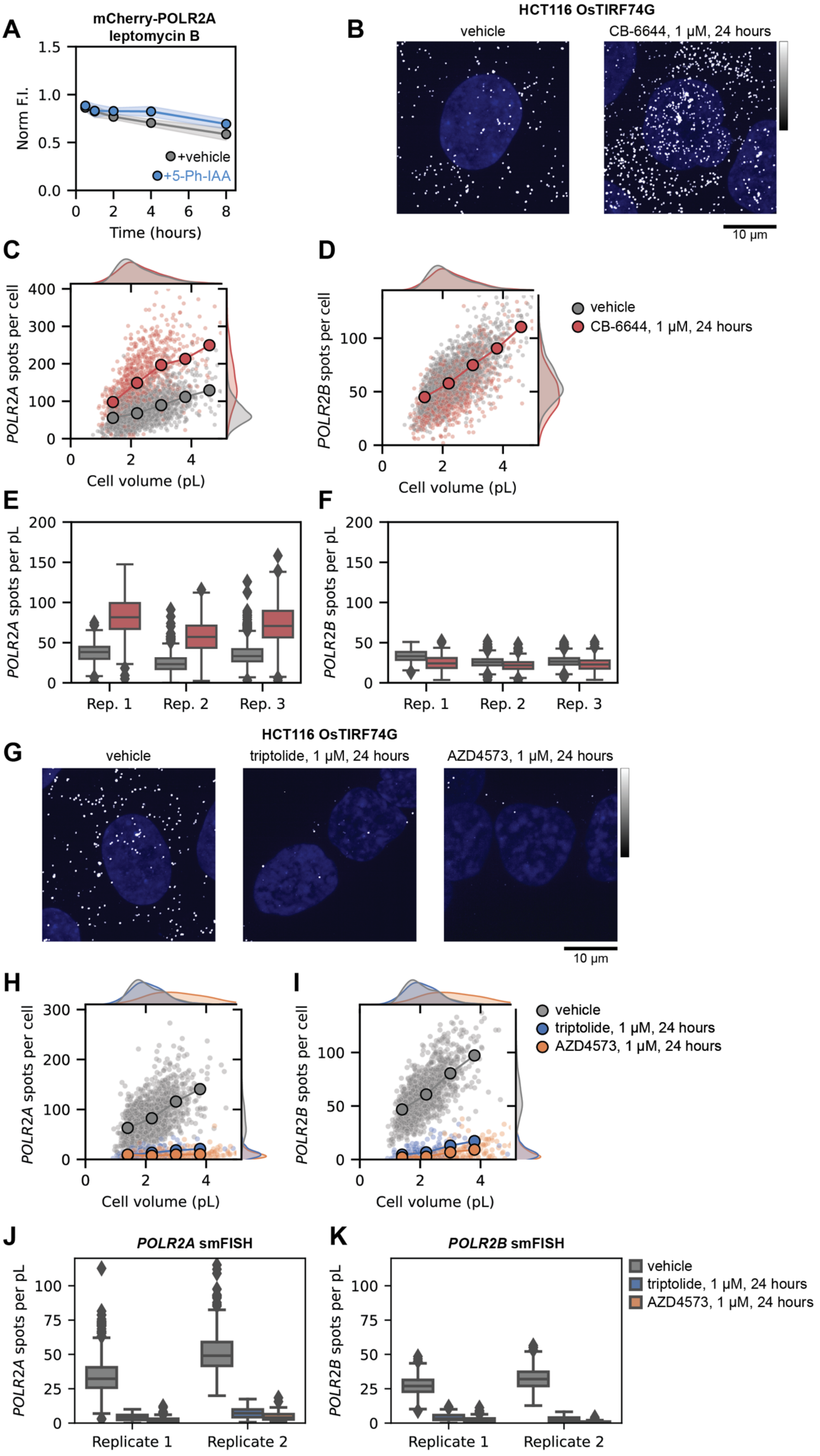
A
mCherry-POLR2A
leptomycin B
Norm F.I.
+vehicle
+5-Ph-IAA
Time (hours)
B
HCT116 OsTIRF74G
vehicle
CB-6644, 1 μM, 24 hours
10 μm
C
POLR2A spots per cell
Cell volume (pL)
D
POLR2B spots per cell
Cell volume (pL)
vehicle
CB-6644, 1 μM, 24 hours
E
POLR2A spots per pL
Rep. 1
Rep. 2
Rep. 3
F
POLR2B spots per pL
Rep. 1
Rep. 2
Rep. 3
G
HCT116 OsTIRF74G
vehicle
triptolide, 1 μM, 24 hours
AZD4573, 1 μM, 24 hours
10 μm
H
POLR2A spots per cell
Cell volume (pL)
I
POLR2B spots per cell
Cell volume (pL)
vehicle
triptolide, 1 μM, 24 hours
AZD4573, 1 μM, 24 hours
J
POLR2A smFISH
POLR2A spots per pL
Replicate 1
Replicate 2
K
POLR2B smFISH
POLR2B spots per pL
Replicate 1
Replicate 2
vehicle
triptolide, 1 μM, 24 hours
AZD4573, 1 μM, 24 hours

**Supplemental Figure 12.**
(A) mCherry-POLR2A after coaddition of 10 ng/µL leptomycin B with 5-Ph-IAA or vehicle in mAC-/mCherry-POLR2A cells.
(B) Example smFISH images of *POLR2A* in HCT116 OsTIRF74G cells treated with vehicle or CB6644 for 24 hours.
(C) smFISH of *POLR2A* transcript count volume-dependence in HCT116 OsTIRF74G cells treated with vehicle or CB6644 for 24 hours.
(D) smFISH of *POLR2B* transcript count volume-dependence in HCT116 OsTIRF74G cells treated with vehicle or CB6644 for 24 hours.
(E) Replicate data for smFISH of *POLR2A* in HCT116 OsTIRF74G cells treated with vehicle or CB6644 for 24 hours.
(F) Replicate data for smFISH of *POLR2B* in HCT116 OsTIRF74G cells treated with vehicle or CB6644 for 24 hours.
(G) Example smFISH images of *POLR2A* in HCT116 OsTIRF74G cells treated with vehicle, triptolide, or AZD4573 for 24 hours.
(H) smFISH of *POLR2A* transcript count volume-dependence in HCT116 OsTIRF74G cells treated with vehicle, triptolide, or AZD4573 for 24 hours.
(I) smFISH of *POLR2B* transcript count volume-dependence in HCT116 OsTIRF74G cells treated with vehicle, triptolide, or AZD4573 for 24 hours.
(J) Replicate data for smFISH of *POLR2A* in HCT116 OsTIRF74G cells treated with vehicle, triptolide, or AZD4573 for 24 hours.
(K) Replicate data for smFISH of *POLR2B* in HCT116 OsTIRF74G cells treated with vehicle, triptolide, or AZD4573 for 24 hours.

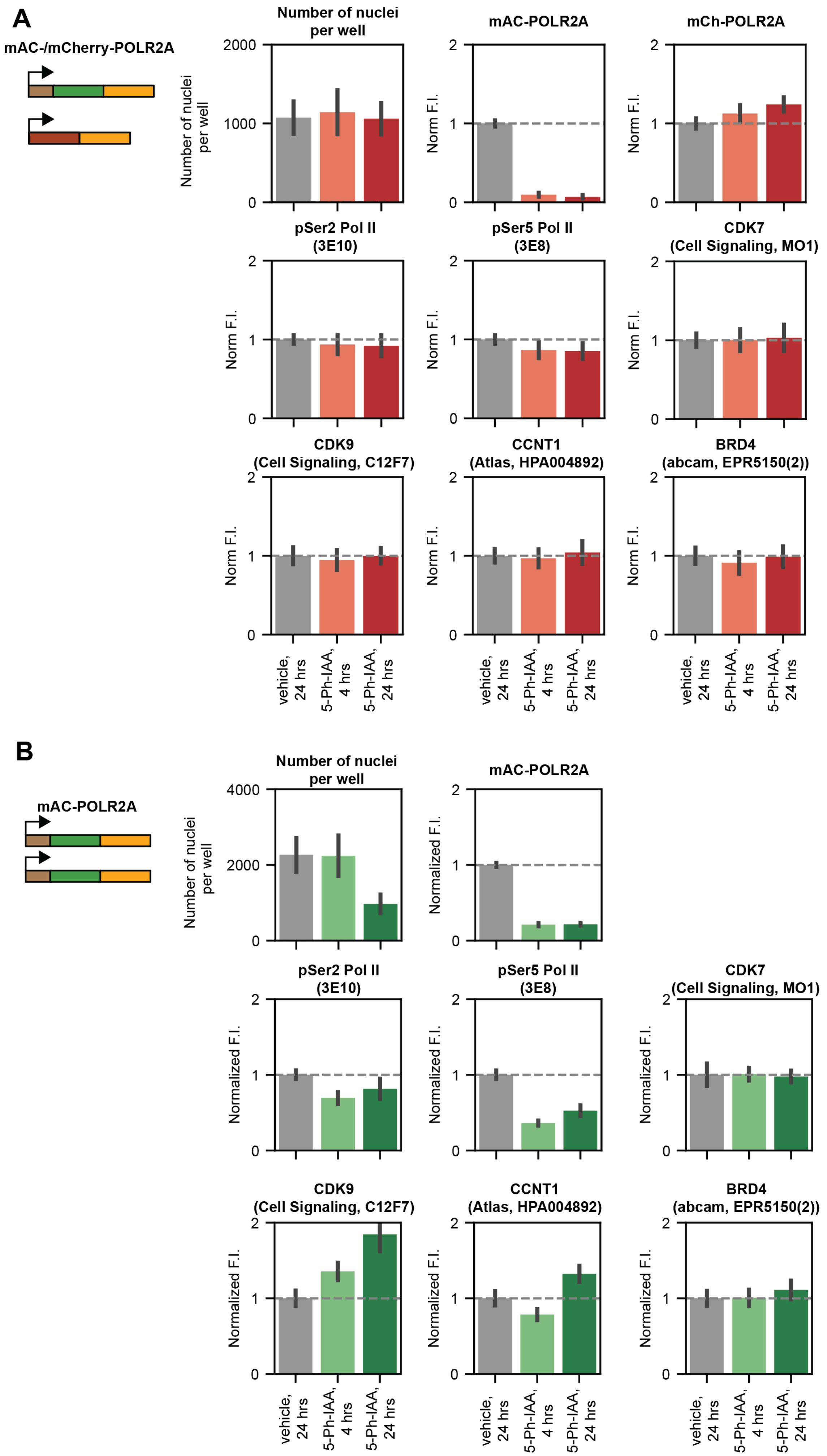
A
mAC-/mCherry-POLR2A
Number of nuclei per well
mAC-POLR2A
mCh-POLR2A
pSer2 Pol II (3E10)
pSer5 Pol II (3E8)
CDK7 (Cell Signaling, MO1)
CDK9 (Cell Signaling, C12F7)
CCNT1 (Atlas, HPA004892)
BRD4 (abcam, EPR5150(2))
Norm F.I.
vehicle, 24 hrs
5-Ph-IAA, 4 hrs
5-Ph-IAA, 24 hrs
B
mAC-POLR2A
Number of nuclei per well
mAC-POLR2A
pSer2 Pol II (3E10)
pSer5 Pol II (3E8)
CDK7 (Cell Signaling, MO1)
CDK9 (Cell Signaling, C12F7)
CCNT1 (Atlas, HPA004892)
BRD4 (abcam, EPR5150(2))
Normalized F.I.
vehicle, 24 hrs
5-Ph-IAA, 4 hrs
5-Ph-IAA, 24 hrs

**Supplemental Figure 13.**

(A) Number of nuclei, mClover-POLR2A and mCherry-POLR2A fluorescence, pSer2 Pol II, pSer5 Pol II, and regulators of Pol II transcription immunofluorescence in mAC-/mCherry-POLR2A cells treated with 5-Ph-IAA for 4 or 24 hours, or vehicle. Bars show mean ± S.D. of replicate wells across two experiments.

(B) Number of nuclei, mClover-POLR2A fluorescence, pSer2 Pol II, pSer5 Pol II, and regulators of Pol II transcription immunofluorescence in mAC-POLR2A cells treated with 5-Ph-IAA for 4 or 24 hours, or vehicle. Bars show mean ± S.D. of replicate wells across two experiments. mAC-POLR2A cells were seeded at twice the density of mAC-/mCherry-POLR2A cells.
